# Vaccination reduces shedding of salmonid alphavirus subtype 3, but bacterial co-infection influences the effect

**DOI:** 10.64898/2026.02.23.707430

**Authors:** Søren Grove, H Craig Morton, Dhamotharan Kannimuthu, HyeongJin Roh, Ravikumar M Chovatia, Ma Michelle Peñaranda, Dawit Ghebretnsae, Kai Ove Skaftnesmo

## Abstract

Waterborne horizontal transmission of viral diseases in fish relies on the release of infectious virus particles (termed shedding) into the aquatic environment. Both the rate and duration of shedding are critical for efficient viral spread, making interventions that reduce shedding valuable for disease control. While vaccines primarily aim to protect individuals from infection and severe disease, they should ideally also limit pathogen transmission by reducing shedding. In this study, we evaluated the capacity of two commercial vaccines - Clynav (DNA vaccine) and AlphaJect Micro 1-PD (inactivated whole-virus vaccine) - to reduce Salmonid alphavirus subtype 3 (SAV3) shedding following experimental infection of Atlantic salmon post-smolts. In individually housed fish, the AlphaJect Micro 1-PD vaccine significantly reduced the proportion of SAV3-shedding fish, the duration of shedding, and the cumulative shedding. The Clynav vaccine significantly reduced the shedding duration and also reduced the proportion of shedding fish. In cohort tanks with concurrent *Tenacibaculum dicentrarchi* co-infection, the AlphaJect Micro 1-PD vaccine significantly reduced the cumulative shedding but increased the number of shedding days. These results demonstrate the potential of vaccines to limit SAV3 transmission, while also highlighting how co-infections likely influence vaccine efficacy.

## 1. Introduction

Aquatic viral diseases pose a major challenge to global aquaculture, where waterborne transmission enables spread both within and between populations. Infected fish release infectious virus particles into the surrounding aquatic environment, a process known as shedding. Shedding has been documented for a number of economically important fish viruses, including Infectious hematopoietic necrosis virus (IHNV) (Mulcahy, Pascho, and Jenes 1983), Virus salmonid herpesvirus-3 (SHV-3) (Faisal et al. 2019), Viral hemorrhagic septicemia virus (VHSV) (Kang et al. 2024), Infectious salmon anaemia virus (ISAV) (Gregory et al. 2009), and Salmonid alphavirus (SAV) (Andersen, Hodneland, and Nylund 2010).

Salmonid alphavirus is an enveloped, positive-sense RNA virus of the family *Togaviridae*. The virus is divided into six or seven subtypes (SAV1-SAV7), based on sequence variation in the E2 and nsP3 genes (Fringuelli et al. 2008; Tighe et al. 2020). Among these, SAV3 and marine SAV2 are particularly associated with pancreas disease (PD) in Atlantic salmon and rainbow trout (*Oncorhynchus mykiss*), causing substantial economic losses in marine aquaculture. Horizontal waterborne transmission of SAV3 is well documented in cohabitation experiments (Graham et al. 2011) and genomic epidemiology studies (Macqueen et al. 2021), and is considered the primary route of PD spread between fish farms (Kristoffersen et al. 2009; Stene et al. 2014; Viljugrein et al. 2009).

The efficiency of waterborne transmission depends on three key factors: the rate and duration of viral shedding; the environmental persistence of viral infectivity; and the minimum infectious dose required for infection. Vaccination aims to induce long-lasting protective immunity by stimulating the adaptive immune system, thereby reducing the risk of infection and severe disease (Plotkin et al. 2018). However, its impact on pathogen transmission is equally important (Chase-Topping et al. 2021; Doeschl-Wilson et al. 2021). So-called ‘leaky’ vaccines, which protect against disease but fail to prevent viral shedding, may paradoxically facilitate pathogen spread (Doeschl-Wilson et al. 2021). Thus, evaluating vaccine effects on shedding is essential for understanding their epidemiological consequences.

Two commercial vaccines are currently available against SAV3. Clynav (Merck Sharp & Dohme Animal Health) is a DNA vaccine encoding the SAV3 structural polyprotein, that is administered intramuscularly. The onset of immunity occurs within 399 degree days, and protection against mortality lasts approximately 9.5 months. AlphaJect Micro1 PD (Pharmaq/Zoetis) is a whole-virus vaccine containing inactivated SAV3 in a liquid paraffin adjuvant, that is administered intraperitoneally. Immunity develops within 516 degree days, with protection lasting ∼12 months. Both vaccines have demonstrated efficacy in reducing mortality and disease severity under controlled laboratory settings (Thorarinsson et al. 2021) and in fish farming conditions (Karlsen et al. 2012; Røsaeg, Thorarinsson, and Aunsmo 2021). Recent field studies have reported superior performance of Clynav in mitigating both mortality and growth retardation during outbreaks involving mixed SAV2 and SAV3 infections (Røsaeg, Thorarinsson, and Aunsmo 2021) and SAV2 alone (Pettersen et al. 2025).

Despite evidence of vaccine efficacy, knowledge regarding their capacity to limit viral shedding, and thus reduce transmission pressure, remains limited. Skjold and coworkers showed that a previously available hepta-valent vaccine (AQUAVAC PD7, MSD) containing five inactivated bacteria and two inactivated whole-viruses including SAV, significantly reduced shedding following intramuscular challenge. Using a cohabitation challenge approach, it has also been demonstrated that the ‘infection pressure’ from vaccinated fish was higher in Micro 1-PD-vaccinated than in Clynav-vaccinated groups (Thorarinsson et al. 2021). Importantly, vaccine effectiveness often varies between controlled laboratory experiments and real-world aquaculture environments (Midtlyng 2016). This discrepancy suggests that vaccine-induced protection is modulated by a range of ‘environmental’ and husbandry-related factors. For example, sea lice treatments and other interventions are known to trigger stress responses in fish and are often associated with increased mortality or disease outbreaks (Overton et al. 2019). Additionally, co-infections with viral (Maj-Paluch et al. 2022), bacterial (Ma et al. 2019; Swaminathan et al. 2021; Li et al. 2025), or parasitic (Bustos et al. 2023) pathogens can influence the host immune response and consequently the degree of vaccine-induced protection (Martins et al. 2011; Figueroa et al. 2017).

RNA viruses have high mutation rates, and a significant proportion the virus released from infected cells and organisms are defective and non-infectious (Duffy 2018). Accordingly, the most accurate and relevant assessment of viral shedding should focus on the subset of particles that are infective. The gold standard for determining viral infectivity involves functional assays, including infection of permissive cell cultures, embryos, or whole organisms to directly measure the capacity of virions to initiate productive infection (Leland and Ginocchio 2007). While these methods provide the most reliable quantification of infective virions, their labour-intensive nature and high costs makes them impractical for large-scale analyses, such as in experiments requiring frequent sampling of multiple replicates. In such contexts, quantification of viral genomes, using RT-qPCR for RNA viruses, offers a valuable and practical proxy. Virus adsorption and elution (VIRADEL) approaches to isolate virus from water (Cliver 1968; Berg, Dahling, and Berman 1971) have been combined with qPCR or RT-qPCR to quantify viral genomes in water samples in numerous studies, including in studies of fish virus shedding (Maheshkumar et al. 1991; Andersen, Hodneland, and Nylund 2010; Bernhardt et al. 2021).

In this study, we compared the effects of two commercial SAV3 vaccines on viral shedding in Atlantic salmon post-smolts following bath challenge. Using a high-throughput VIRADEL and RT-qPCR approach, we analysed 1,233 water samples from 39 fish tanks over 32-35 days post-challenge. This approach enabled a robust comparison of vaccine effects on transmission potential, addressing a critical gap in understanding the epidemiological consequences of vaccination against SAV.

## 2. Material and Methods

### 2.1 Fish and vaccination

Atlantic salmon were hatched and reared at the Institute of Marine Research (IMR) Matre Research Station (Matre, Norway) following standard protocols. Broodfish originated from wild Atlantic salmon from the river Etne. Prior to vaccination, smoltification was induced using a 12-hour light:12-hour darkness (‘winter signal’) for 9 weeks. Three distinct groups of salmon were established for the experiment (**Figure 1**). The first group, consisting of 283 individuals, was vaccinated with Clynav (MSD Animal Health) in accordance with the manufacturer’s guidelines. The second group, also comprising 283 individuals, was vaccinated with AlphaJect Micro1-PD (Pharmaq AS), following the manufacturer’s instructions. A third group of 423 (i.e. 140 + 283) salmon parr was maintained as an unvaccinated control. To account for potential losses prior to the experiment, 40 additional fish per group were included in each group but are not shown in **Figure 1**. Post-vaccination, all fish were kept under continuous light conditions until their transfer. While still in freshwater, the fish groups were transferred to the IMR Disease Laboratory in Bergen (Norway). Upon arrival, the fish were acclimated to 20 ppt seawater during a 24-hour period, after which the salinity was increased to 35 ppt seawater. During the remaining experimental period, fish were kept at a 12:12hr light regime, in 35 ppt seawater at 10-12°C. The fish were fed once a day with Nutra Sprint 3.0 mm (Skretting). Prior to the start of experiments, 20 unvaccinated salmon were tested for viral infections by TaqMan RT-qPCR assays and found negative for the presence of salmonid alphavirus subtype 3 (SAV3) (Hodneland and Endresen 2006), infectious pancreatic necrosis virus (IPNV) (Nylund et al. 2011), piscine myocarditis virus (PMCV) (Lovoll et al. 2010), and piscine orthoreovirus-1 (PRV-1) (Wessel et al. 2015). Fish were anaesthetized using 100 mg/L tricaine methanesulfonate (Finquel MS-222, MSD Animal Health) before handling and euthanized with an overdose of Benzocaine (200 mg/L) before sampling. All experimental procedures were reviewed and approved by the Norwegian animal research authority prior to the start of challenge experiments (FOTS ID: 31161).

**Figure 1.**
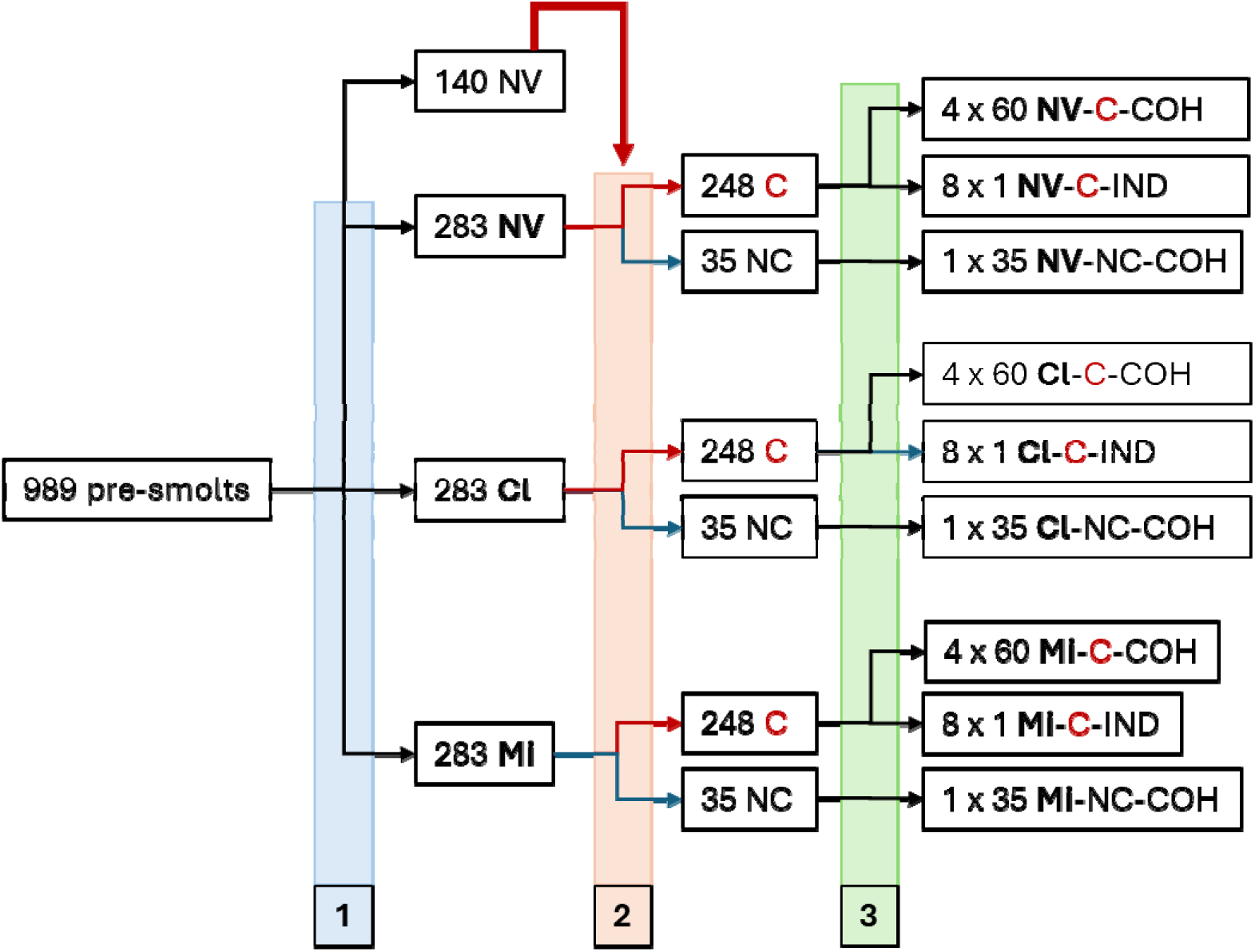
Overview of experimental design and treatment groups. A group of 989 Atlantic salmon pre-smolts were divided into four groups. (1) One group of 140 fish were left unvaccinated and kept for later use as shedder fish. Other 283 fish were left as an unvaccinated (NV) group, while two groups of 283 fish were vaccinated with Clynav (Cl) and AlphaJect Micro1-PD (Mi), respectively. (2) Eight weeks post-vaccination, and after smoltification and transfer to seawater, each of the NV-, Cl- and Mi-groups were divided into 35 fish that were left unchallenged (NC) and 248 fish that were bath challenged (C) with water from the 140 shedder fish. (3) After bath challenge, each challenged group was divided between 4 cohort tanks (COH) of 60 fish, and 8 individual tanks (IND) with one fish each.

### 2.2 Bath challenge

Sixty-one days (∼370 degree days) after vaccination, a bath challenge of vaccinated and unvaccinated groups was performed. The average weight of the challenged salmon was 61.8±9.4 g. Ten days prior to the bath challenge, a total of 140 salmon smolts from the unvaccinated group received an intraperitoneal injection of 100 μL cultured SAV3 (5×10^4^ TCID_50_/mL in L15 cell culture medium), corresponding to 5×10^3^ TCID_50_/fish. The 140 ip. injected fish, hereafter called ‘bath challenge shedders’ were kept in a 400 L tank supplied with 35 ppt seawater at a flow rate of 600 L/hour. Ten days after the ip. injection, the water flow was stopped, and the bath challenge shedder fish were kept with ample aeration for 8 hours to shed infectious SAV3 into the water. Water samples were collected and filtered as described below (section 2.3) at 0-, 1-, 2-, 3-, 4-, 5-, 6-, 7- and 8-hours after stoppage of the water flow. After 8 hours in static water, the 140 bath challenge shedders were transferred to new tanks, and the water remaining in the tank used to bath challenge vaccinated and unvaccinated groups of salmon smolts as follows. In three 400 L tanks, containing either 300 Clynav vaccinated, 300 AlphaJect Micro1-PD vaccinated or 300 unvaccinated fish, respectively, the water volume was very briefly lowered to approximately 140 L. 110 L of water from the bath challenge shedder tank was immediately added to each of the tanks. The tanks were then left with the water flow stopped for 6 hours, with ample aeration and mechanical pumps supplying water circulation. After 6 hours, the bath challenge was stopped by turning on the water flow at 600 L/hour. After one hour, 283 challenged fish from each group were netted into buckets with clean seawater and transferred to the tanks from where water would be sampled during the rest of the experiment (**Figure 1**). Each group of challenged fish was divided between 4 cohort tanks (60 fish per 250 L tank) and 8 individual tanks (one fish per 20 L tank). Three 150 L tanks containing either 35 unchallenged Clynav vaccinated, 35 unchallenged AlphaJect Micro1-PD vaccinated or 35 unchallenged unvaccinated fish, respectively, were kept as negative controls. Surplus fish, i.e. ∼20 Clynav vaccinated, ∼20 AlphaJect Micro1-PD vaccinated and ∼20 unvaccinated fish, remained in the tanks in which they were challenged.

### 2.3 Water sampling and filtration

Water samples of 100 mL were filtered directly from the fish tanks following the time schedule specified in **Table 1**. Prior to sampling from a tank, the water supply was stopped for exactly three hours. Each water sample was collected from the tanks using a peristaltic pump, a Watson Marlow 520S for the cohort tanks and a Heidolph Hei-Flow Value 1 for the individual tanks, and directly pumped through a 1.0 μm Nucleopore prefilter (φ 25 mm, Cytiva/Whatman 10418718) followed by a 0.45 μm Cellulose Nitrate filter (φ 13 mm, Cytiva/Whatman 7184-001). The Nucleopore prefilter was mounted in a Swinnex 25 mm filter holder (Merck, SX0002500) and the Cellulose Nitrate filter in a Swinnex 13 mm filter holder (Merck, SX0001300). After removal from the filter holder, each 13 mm Cellulose Nitrate filter was briefly placed on a paper tissue to remove excess water, then folded twice, and transferred to a homogenization tube (PreMax Matrix 96 tubes 2D, LVL Technologies) containing two steel beads, and immediately frozen at −20°C. For long term storage, the filters were stored at – 80°C.

**Table 1.** Overview of the time points (dpc) where water samples were collected and filtrated from cohort and individual tanks, respectively. S, sampling performed; NS, sampling not performed. 1) Weekend days.

| Tanks | Sampling (dpc) |  |  |  |  |  |  |  |  |  |
| --- | --- | --- | --- | --- | --- | --- | --- | --- | --- | --- |
|  | 2-4 | 5-6<sup>1)</sup> | 7-11 | 12-13<sup>1)</sup> | 14-18 | 19-20<sup>1)</sup> | 21-25 | 26-27<sup>1)</sup> | 28-32 | 33-34<sup>1)</sup> |
| Cohort | S | S | S | S | S | S | S | S | S | NS |
| Individual | S | NS | S | S | S | S | S | S | S | S |

### 2.4 Nucleic acid extraction from Cellulose Nitrate filters

The filters were thawed at room temperature, 500 μL homogenization buffer (Lybe Scientific LSNXE0384) was added to each tube, and the filters then homogenized using a Spex 1600 MiniG tissue homogenizer (SPEX SamplePrep). The homogenized filters were spun down by centrifugation at 1100 *g* for 5 minutes in an Allegra X-30R centrifuge (Beckman Coulter), and nucleic acids were extracted from aliquots of 200 μL of supernatant using the NAxtra 2.0 Aquaculture NA extraction kit (Lybe Scientific LSNXE0384) on a Biomek 4000 Automated Laboratory Workstation (Beckman Coulter). In the final step of the extraction process, the nucleic acids were eluted in 50 μL of elution buffer (Lybe Scientific).

### 2.5 RT-qPCR analysis of SAV3 RNA levels in water samples (Cellulose Nitrate filters)

Nucleic acid extracts were diluted tenfold in RNase-free water. For each reaction, 4 μL of the diluted extract was used as the template in a 10 μL total volume RT-qPCR reaction, performed using the AgPath-ID One Step RT-PCR kit (Thermo Fisher Scientific). Primers and probes targeted the nSP1 region of SAV3 as previously described (Hodneland and Endresen 2006; Kannimuthu et al. 2026). Thermocycling was conducted on a 384-well Quantstudio5 qPCR instrument (Thermo Fisher Scientific). Briefly, RT-qPCR conditions consisted of a reverse transcriptase incubation step at 45°C for 10 minutes and a polymerase activation step at 95°C for 10 minutes followed by 40 cycles of denaturation at 95°C for 15 seconds and a combined anneal and extension step at 60°C for 45 seconds.

### 2.6 Quantitative PCR (qPCR) analysis of three *Tenacibaculum* species in water samples (Cellulose Nitrate filters)

During the experiment, all fish in the cohort tanks developed clinical signs of a bacterial infection, which were later attributed to *Tenacibaculum* spp. Thus, further diagnostic testing by qPCR was conducted to identify the specific bacterial species. Primers and probes for *Tenacibaculum dicentrarchi*, *Tenacibaculum maritimum* and *Tenacibaculum finnmarkense* were designed with Speciesprimer (Dreier et al. 2020). Ten μL multiplex qPCR reactions using TaqMan™ Multiplex Master Mix (Thermo Fisher) with 300 nM primer concentration for each primer and 250 nM of each probe and 4 μL of 1:10 diluted total nucleic acid extracts from filters were prepared. Thermocycling was conducted on a Quantstudio 5 instrument (Thermo Fisher) using an initial denaturation step at 95°C for 20 seconds and 40 cycles of denaturation at 95 for 1 second and a combined annealing/extension step at 60°C for 20 seconds. Primers and probe sequences are listed in supplementary information (**Supplementary Table 1**).

### 2.7 Tissue sampling

At 14 dpc, eight fish from each challenged group were sampled from the surplus fish tanks. At 32 dpc, eight fish were sampled from each cohort tank, which amounted to eight fish from each negative control group and 32 fish from each challenged group. At 35 dpc, all the fish from the single-fish tanks were sampled. At each sampling, the fish were euthanised by an overdose of 200 mg/L tricaine methanesulfonate (Finquel MS-222) and then heart, pancreas, spleen and skeletal muscle tissue dissected. Skeletal muscle was dissected from the lateral side under the dorsal fin and included the sideline; and the heart was cut in two halves. One half of the heart, spleen, and a part of the pancreas were transferred to separate sampling tubes (PreMax Matrix 96 tubes 2D, LVL Technologies) containing 600 µL homogenization solution (Promega Maxwell HT simplyRNA kit, Custom), and kept on ice until homogenization using a bead-based tissue homogenizer (SPEX SamplePrep 1600 MiniG) at 1300 rpm for 5 mins. Homogenates were stored at −80°C until RNA extraction. The other half of the heart, skeletal muscle, and the remaining part of the pancreas tissue were transferred to 10% buffered formalin and stored for 48 hours at room temperature and then transferred to 70% ethanol for long-term storage.

During the experiment, samples of skin and skin lesions were collected from moribund and recently dead cohort fish and transferred to 10% buffered formalin as described above.

### 2.8 RNA extraction from tissue samples, and RT-qPCR analysis of extracted RNA

Total RNA was extracted from heart, pancreas, spleen and skeletal muscle tissues using Promega Reliaprep simplyRNA HT 384 (Nerliens) on a Biomek 4000 Laboratory Automated Workstation according to the manufacturer’s instructions and quantified using a NanoDrop-1000 spectrophotometer (Thermo Fisher Scientific). The RNA samples were normalized to a concentration of 50 ng/µL using the Biomek 4000 Laboratory Automated Workstation. RT-qPCR for SAV3 was performed using 2 μL of template as described in Section 2.5 (Kannimuthu et al. 2026).

### 2.9 Histopathology and immunohistochemistry

Heart, skeletal muscle, and pancreas samples in 70% ethanol were processed using a tissue processor (Leica TP1020) and embedded in Histowax (HistoLab). Paraffin blocks were sectioned at 3 µm thickness using a rotary microtome (Thermo Scientific HM355S). Sections were stained with hematoxylin, erythrosine, and saffron (HES). After mounting with coverslips using Histokitt (Chemi–Teknik AS), slides were scanned at 40× magnification using a NanoZoomer S60 digital slide scanner (Hamamatsu Photonics) and examined with NDP.view2 software (Hamamatsu Photonics). Lesions were scored semi-quantitatively for inflammation and degeneration or necrosis as previously described (McLoughlin et al. 2006; Finstad et al. 2012; Herath et al. 2017) (**Supplementary Table 2**).

Sections of skin lesions from moribund cohort fish were analyzed using modified Gram staining and immunohistochemistry (IHC) to investigate the etiology of the lesions. The modified Gram staining followed the Sigma–Aldrich kit (Sigma-Aldrich, cat#1.01603.0001) protocol with a key modification: after staining with fuchsin, sections were incubated in a 0.74% formaldehyde and 1% glacial acetic acid solution to enhance visualization of Gram-negative bacteria. Following this, sections were counterstained with saffron for 5 minutes, washed in tap water, and dehydrated through graded alcohol. Finally, sections were then cleared in xylene and mounted with coverslips.

For immunohistochemistry, antigen retrieval was performed in citrate buffer (pH 6.0) using an Antigen Retriever 2100 (Aptum Biologics Ltd.) for 20 minutes. Endogenous alkaline phosphatase activity was inhibited by incubating sections with a dual endogenous enzyme block reagent (Dako, S2003) for 10 minutes, and non-specific binding was blocked by incubation in 5% bovine serum albumin (BSA) in 1X Tris buffer for 20 minutes. Between steps, sections were washed for 5 minutes in 1X Tris buffer containing 1% Tween-20.

Sections were incubated overnight at 4 °C with a polyclonal rabbit anti-*Tenacibaculum* spp. primary antibody (Sandlund et al. 2021) diluted 1:500 in 2.5% BSA in 1X Tris buffer. Following primary antibody incubation, sections were incubated with a biotinylated horse anti-mouse/rabbit IgG secondary antibody (Vector Laboratories, BA-1400) for 20 minutes, followed by incubation with VECTASTAIN ABC–AP reagent (Vector Laboratories, AK-5001) for 30 minutes at room temperature. Immunoreactivity was visualized using the ImmPACT Vector Red AP substrate kit (Vector Laboratories, SK-5105) for 20–30 minutes. Finally, washed sections were rinsed in tap water, counterstained with instant hematoxylin for 1 minute, dehydrated in absolute alcohol, cleared in xylene, and mounted with Histokitt (Chemi-Teknik AS).

### 2.10 Statistical analysis

Statistical analysis included the following tests, performed in GraphPad Prism 10: nonparametric Kruskal-Walli’s test (Kruskal and Wallis 1952), Šidák’s multiple comparison test (Šidák 1967), two-way ANOVA and Tukey’s multiple comparison test (Tukey 1949), Dunnett’s multiple comparison test (Dunnett 1955), Dunn’s nonparametric multiple comparison test (Dunn 1961), Fisher’s exact test (Fisher 1922), and Spearman’s rank correlation test (Spearman 1904).

## 3. Results

### 3.1 Experimental observations

#### Determination of optimal water sampling parameters

To prepare infectious water for the bath challenge, the water inflow to a tank containing SAV3-infected shedder fish was stopped for eight hours. Water samples were collected from the tank every hour during the stoppage and analysed for SAV3 RNA levels by RT-qPCR (**Figure 2**). The results demonstrated that the SAV3 RNA concentration increased markedly during the initial 3 hours, followed by a slower accumulation. Over the 8-hour period, the mean Ct value decreased by ΔCt = 4.39, indicating an >20-fold increase in viral RNA. The time required to filter 100 mL water samples increased progressively with time after inflow stoppage, reaching ∼5 minutes at 3 hours – a stoppage time point associated with a ∼10-fold (ΔCt = 3.25) increase in SAV3 RNA levels (**Figure 2**). We found that, particulate material (mucus, scales, faeces) accumulated in the tank water after the flow was stopped, and progressively clogged the filters and increased the filtration time. A 3-hour inflow halt was accordingly adopted in the standard protocol for all subsequent water sampling (section 2.3).

**Figure 2.**
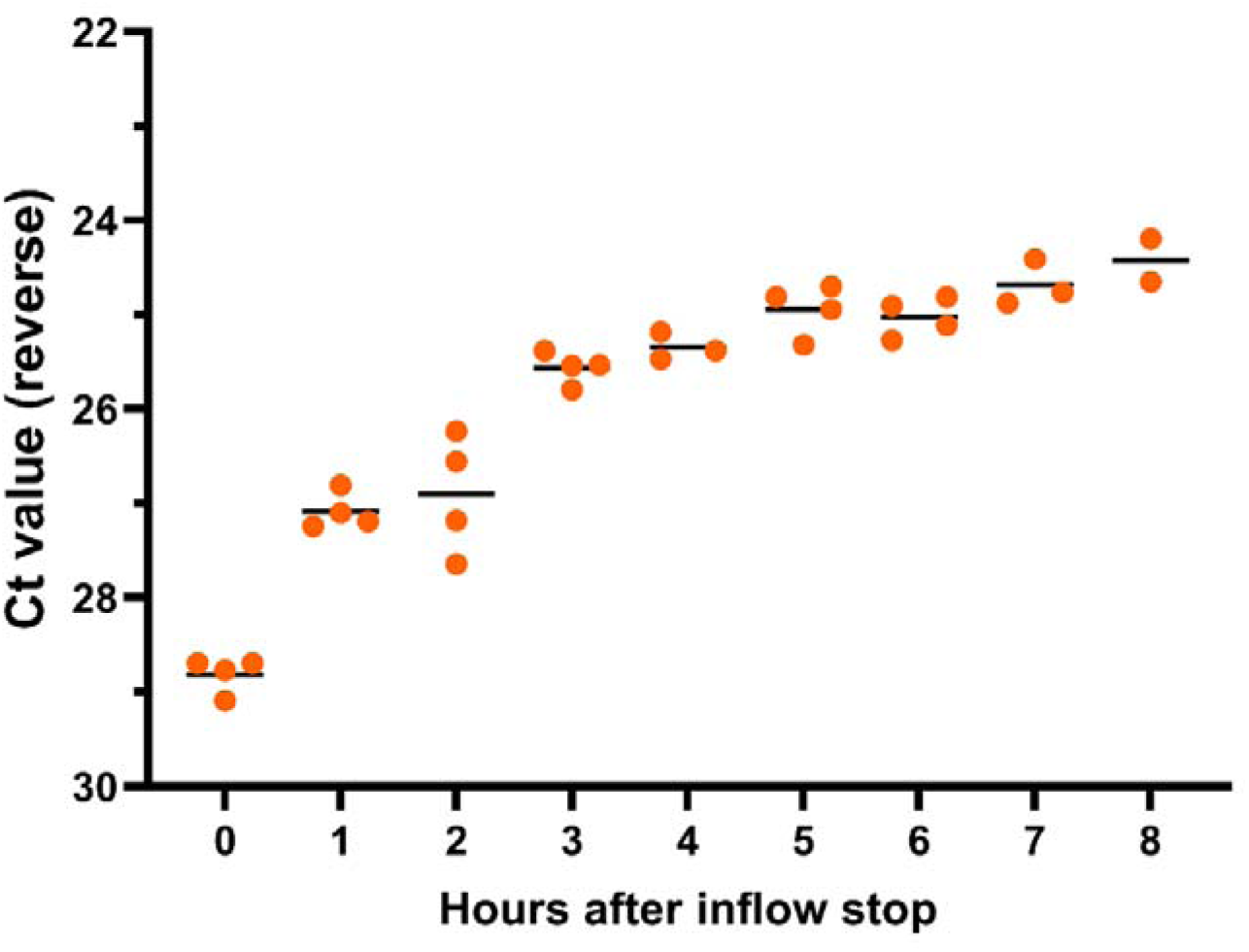
SAV3 RNA levels in water samples collected hourly from a shedder fish tank. The water supply to a shedder fish tank was stopped for eight hours before water from the tank was used to bath challenge groups of non-vaccinated, Clynav-vaccinated and AlphaJect Micro1-PD vaccinated salmon post-smolts, respectively. Water samples were taken every hour after the stoppage and analyzed by RT-qPCR for their content of SAV3 RNA. Each point represents an independent water sample, and horizontal lines show the mean Ct value of samples taken at each time point. The y-axis represents reverse Ct values, and the x-axis shows hours after inflow stoppage.

#### Morbidity and mortality

Following the SAV3 bath challenge and transfer of fish to their final, individual or cohort tanks, a small number of fish were euthanized due to erratic behaviour during the first 3 days post-challenge across all cohort groups (**Figure 3**). Most of these fish exhibited no macroscopical clinical signs, except for one fish in the Cl-C cohort which displayed small ulcers along the lateral line and around the dorsal fin. Between 4 and ∼20 dpc, moribund and dead fish were observed at a low frequency in all cohort tanks. After 18-20 dpc, cumulative mortality increased in the NV-NC and Cl-NC control tanks. From approximately 27-28 dpc, elevated mortality was also observed in some of the challenged cohort tanks (**Figure 3**). Due to increasing mortality rates, the cohort experiment was terminated at 32 dpc as a humane endpoint. At this termination, 119 of 120 sampled fish across all cohorts exhibited varying degrees of the macroscopical clinical signs in fins described for moribund and dead fish below. In contrast to the cohort experiment, no mortality was observed in fish housed in individual tanks, and all the fish were visually and macroscopically in good condition when this experiment was terminated at 35 dpc. The SAV isolate used in this experiment have not caused mortality in challenged salmon post-smolts in previous experiments (Roh et al. 2024; Kannimuthu et al. 2026).

**Figure 3.**
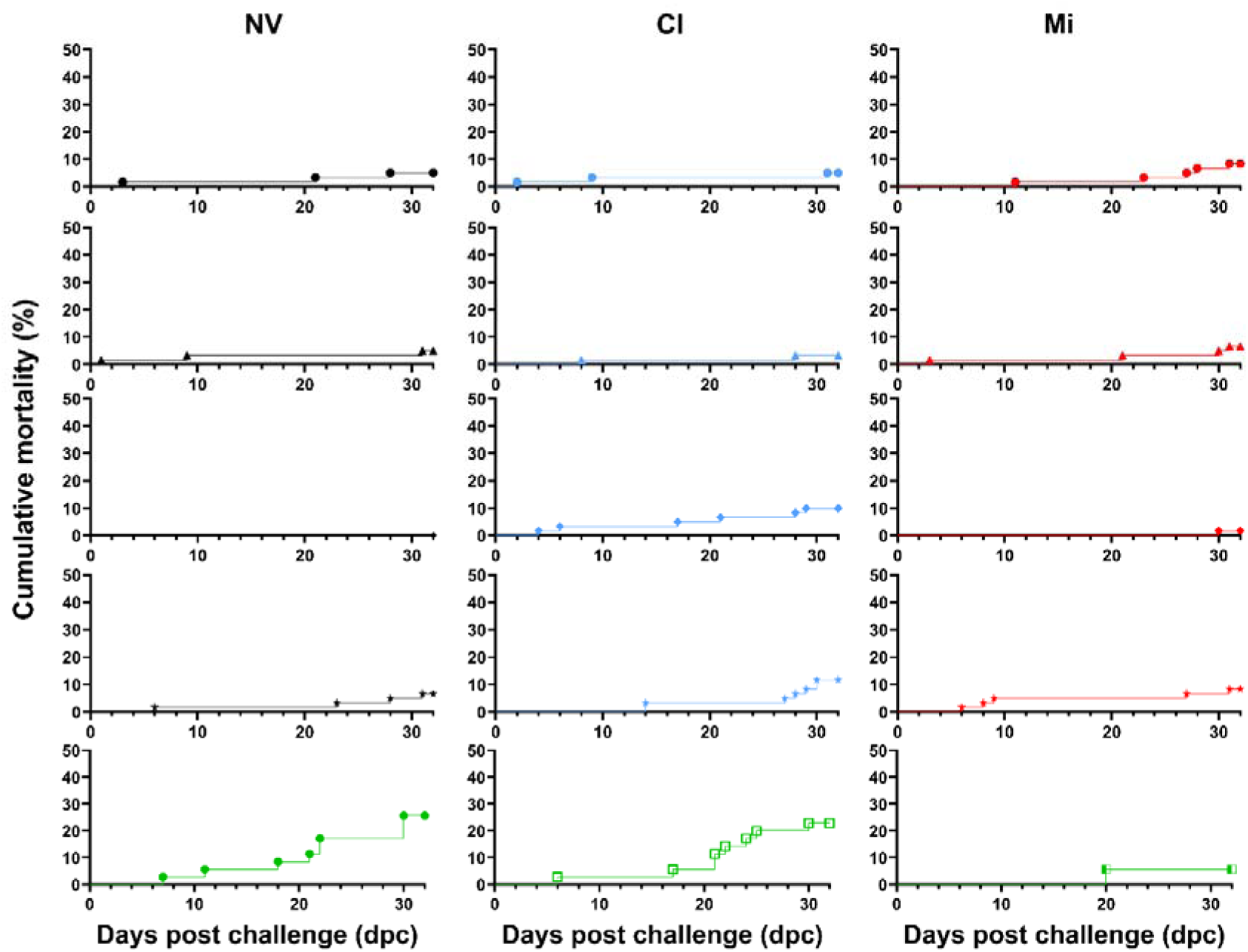
The cumulative mortality in SAV3 challenged and non-challenged cohort tanks. Mortalities, including the humane endpoint of moribund fish, were recorded for all cohort tanks. For each vaccination group, the four upper panels display the results from four challenged replicate tanks (NV in black, Clynav in blue and AlphaJect Micro1-PD in red) and the lower panel shows results from a corresponding non-challenged tank (green). The y-axis represents the cumulative mortality in percent, and the x-axis shows days post challenge (dpc).

#### Bacterial co-infection

Moribund fish and dead fish observed during the cohort experiment typically exhibited loss of scales, substantial fin erosion and open ulcers, occasionally with a yellowish wound edge (**Supplementary Figure 1**). Euthanized moribund fish were sampled for histopathological, immunohistochemical and qPCR analyses. Microscopy analysis of tissue sections stained with HES demonstrated the presence of widespread colonies of basophilic, long rods or filamentous–like bacteria across much of the epidermis. Affected regions exhibited significant inflammation, necrosis, and bleeding. In the most severe cases, the infection caused deep lesions that completely eroded the skin layers, including the epidermis, dermis, and hypodermis, as well as the underlying red muscle. Furthermore, bacteria were found to have penetrated deep into the white muscle tissue. Modified Gram staining of skin lesion sections revealed gram-negative rod-shaped, filamentous bacteria consistent with *Tenacibaculum* spp (**Supplementary Figure 2**). Immunohistochemical staining of skin lesion sections using a polyclonal antiserum against *Tenacibaculum* confirmed the abundant presence of *Tenacibaculum* spp. in these ulcerated regions (**Supplementary Figure 3**). Analysis by qPCR demonstrated only *Tenacibaculum dicentrarchi* in ulcers, with no evidence of *Tenacibaculum finnmarkense* or *Tenacibaculum maritinum*. Additionally, filters from the water samplings aimed at detecting shedding of SAV3 were analysed for *T. dicentrarchi*, *T. finnmarkense* and *T. maritinum* by qPCR. Despite that the filtration protocol was optimized for SAV3 RNA detection (including a 1.0 μm prefilter), DNA from *T. dicentrarchi* was detected in water from all cohort tanks (**Supplementary Figure 4**). Neither *T. finnmarkense* nor *T. maritinum* were detected in these water samples. Despite absence of signs of *Tenacibaculum* presence among the individually housed fish, *Tenacibaculum dicentrarchi* was sporadically detected in water samples from fish from all vaccination groups (**Supplementary Figure 5**).

### 3.2 Fish weight at experiment termination

At experiment termination, the weight of all individually housed fish were recorded (35 dpc), as where the weights of eight fish sampled from each cohort tank at 32 dpc. For each vaccination group (NV, Clynav and Micro1-PD), the mean weight of the individually housed fish were compared to the mean weights of fish from the corresponding cohort tanks (**Figure 4**). For all vaccination groups, the mean weight of individually housed fish was higher than that of fish from both challenged and non-challenged cohort tanks. While these differences were not statistically significant for the NV group (Dunnett’s multiple comparison test), individually housed fish in the Clynav and Micro1-PD groups had significantly higher mean weight compared to their challenged cohort counterparts. Additionally, in the Micro1-PD group, the mean weight of individually housed fish was also significantly higher than that of the non-challenged tank (NC) (**Figure 4).** No significant differences in mean weight were observed between the individually housed vaccination groups (NV-C, Cl-C and Mi-C) (**Figure 5A**).

**Figure 4.**
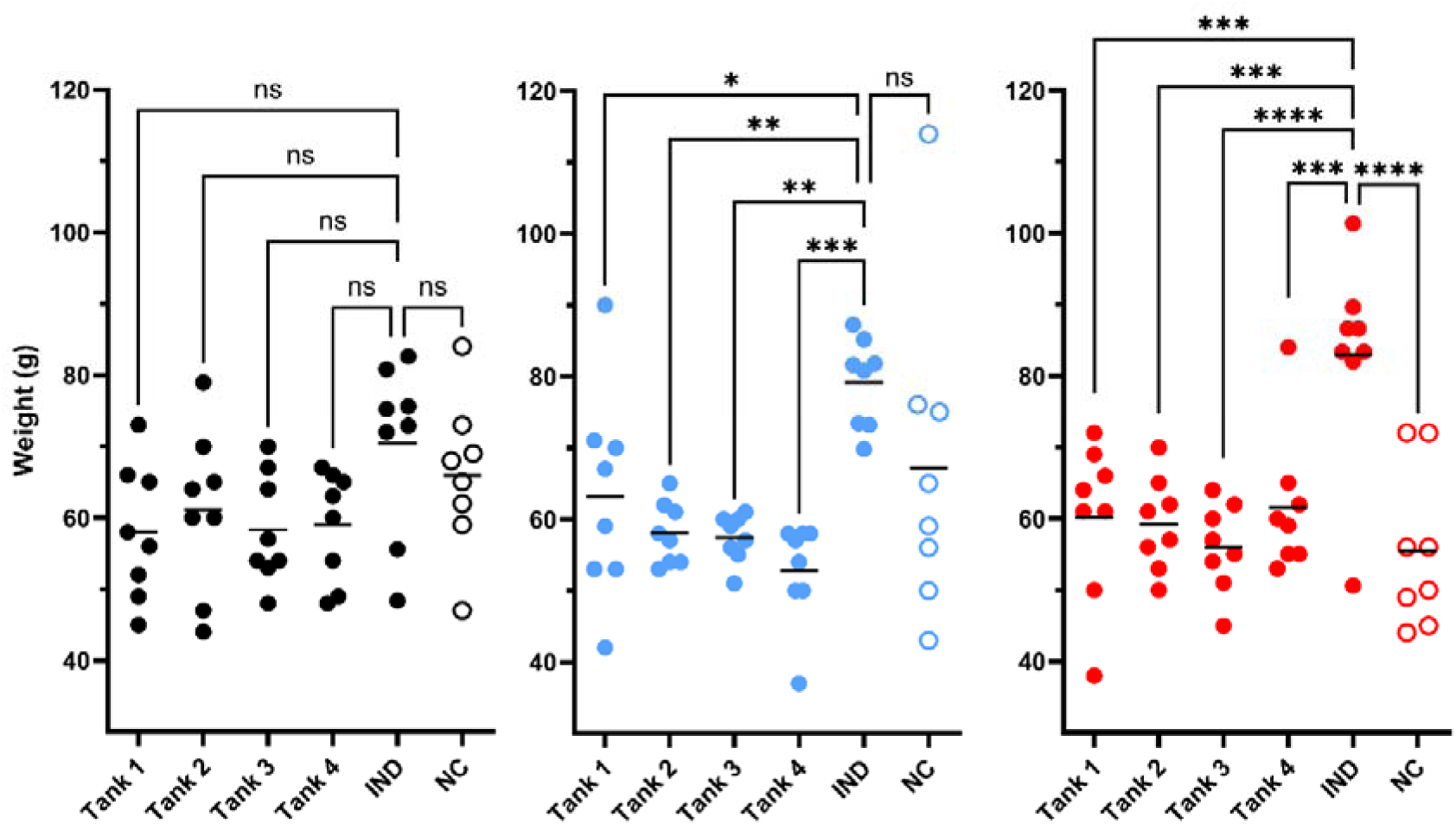
Weight of sampled cohort fish and individual fish at experiment termination. Eight fish from each cohort tank were sampled at 32 dpc, and 8 fish from each individual group were sampled at 35 dpc. Individual points represent the weight of each fish, showing the spread and variability within each group. Dots represent SAV3 challenged groups, and circles represent non-challenged groups (NC). Black color: non-vaccinated; blue color: Clynav vaccinated; red color: AlphaJect Micro1-PD vaccinated. Tank 1-4: four replicate cohort tanks, IND: fish from individual tanks. The y-axis represents fish weight (g), and horizontal bars show the mean weight of the groups. Asterisks (ns, not significant; * *p* < 0.05; ** *p* < 0.01; *** *p* < 0.001: **** p < 0.0001) indicate statistically significant differences between cohort vaccination groups and the corresponding individual group (IND) based on a Dunnett’s multiple comparison test (one-way ANOVA).

**Figure 5.**
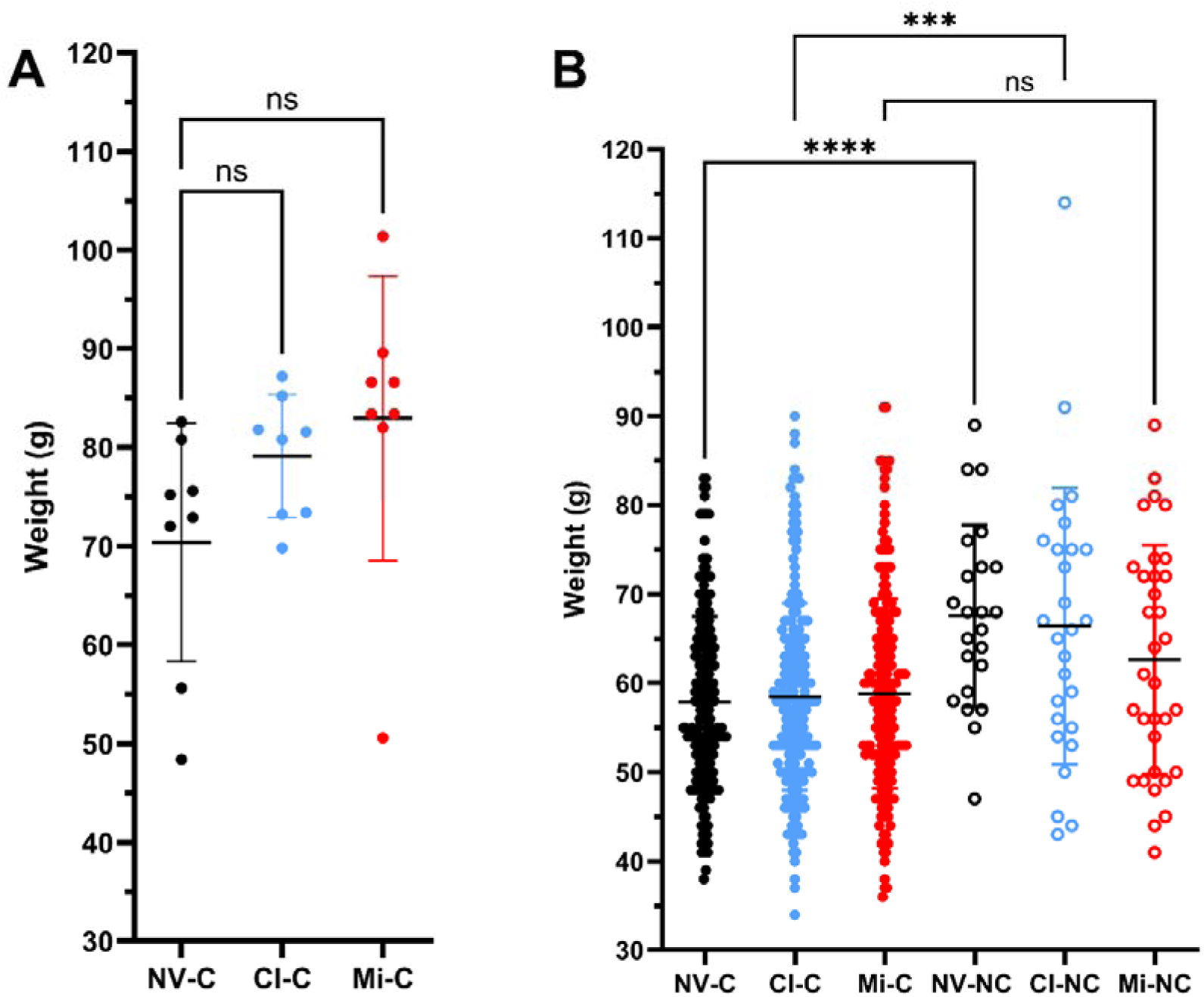
Weight of individual fish and surviving cohort fish at experiment termination. All eight fish from each individual group were sampled at 35 dpc and their weight recorded (A). In the cohort experiment, eight fish from each cohort tank were sampled and weighed at 32 dpc, and all remaining fish euthanized and weighed at 35 dpc (B). Individual points represent the weight of each fish, showing the spread and variability within each group. Black color: non-vaccinated (NV); blue color: Clynav vaccinated (Cl); red color: AlphaJect Micro1-PD vaccinated (Mi). Dots represent SAV3 challenged groups (C), and circles represent non-challenged groups (NC). The y-axis represents fish weight (g), and horizontal bars show the mean weight of the groups. In (A), ns indicates not significant by Dunnett’s multiple comparison test (one-way ANOVA). In (B), ns indicates not significant and *** indicates p < 0.001 by Šidák’s multiple comparison test (one-way ANOVA).

In the cohort experiment, fish not sampled at 32 dpc, were euthanized and weighed at 35 dpc. When including the weights of all cohort fish, measured at either 32 or 35 dpc, the mean weights of challenged NV-vaccinated (NV-C) and Clynav-vaccinated (Cl-C) groups were significantly lower (Šidák’s multiple comparison test) than that of the corresponding non-challenged groups (NV-NC and Cl-NC, respectively) (**Figure 5B**). Although the mean weight of the challenged Micro1-PD-vaccinated group (Mi-C) was lower than that of the non-challenged group (Mi-NC), this difference was not significant. No significant differences in mean weight were seen among the challenged vaccination groups or among non-challenged vaccination groups (data not shown).

### 3.3 Kinetics of SAV3 shedding

Water analysis was used to monitor the kinetics of SAV3 shedding in both individually housed and cohort-based experiments. This approach allowed for the quantification and comparison of viral RNA shedding across different vaccination groups, providing insights into the efficacy of each vaccine in limiting SAV3 transmission.

For individually housed fish, the results showed that all eight of eight challenged non-vaccinated (NV-C) fish shed SAV3 RNA to the water, compared to four of eight Clynav-vaccinated (Cl-C) fish and none of eight Micro1-PD-vaccinted (Mi-C) fish (**Figure 6**). By Fisher’s exact test, the number of fish shedding in the Mi-C group was significantly lower than the number in the NV-C group (p=0.0002), while the difference between the Cl-C and NV-C groups was not significant (p=0.0762). The last day where shedding was detected at 16 dpc in the NV-C group and at 15 dpc in the Cl-C group. A highly significant association (Fisher’s exact test, p<0.00001) was observed between occurrence of SAV3 shedding and the presence of SAV3 RNA in heart tissue at 35 dpc. Specifically, 11 fish both shed virus and tested positive in the heart, one fish shed virus but was heart-negative, one fish did not shed virus but was heart-positive, and 13 fish neither shed virus nor tested positive in the heart (**Figure 6**). Furthermore, the duration of viral shedding, measured as the number of days shedding was detected, differed significantly between the NV-C group and both the Cl-C (Tukey’s multiple comparison test, p=0.0012) and Mi-C (p<0.0001) groups, (**Figure 7A).**

**Figure 6.**
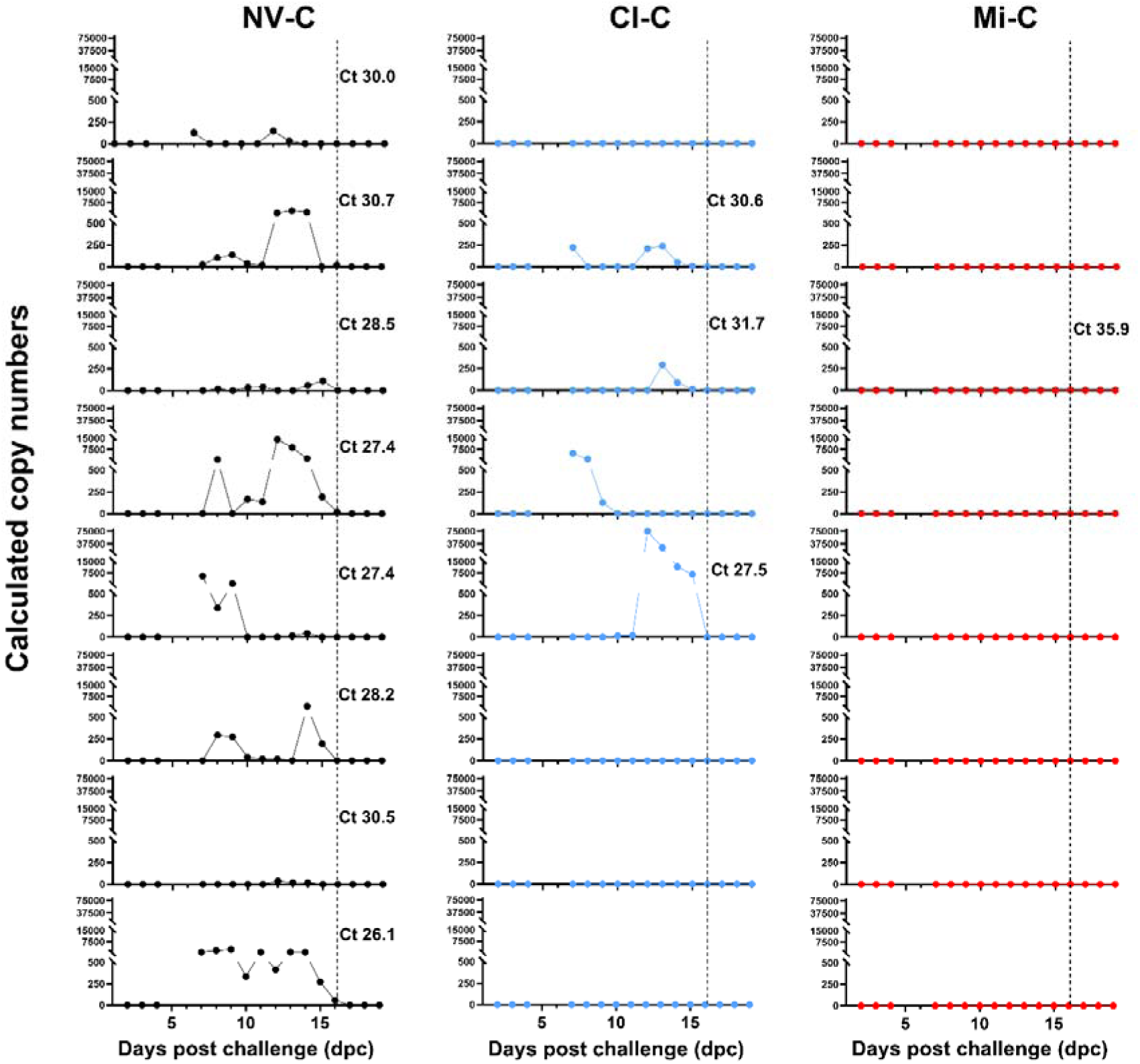
Temporal virus shedding profiles in individual fish after SAV3 bath challenge. Water samples (100 mL) were collected daily from eight individual fish per challenged vaccination group (Non-vaccinated, NV-C; Clynav, Cl-C and AlphaJect Micro1-PD, Mi-C) between 2- and 35-days post-challenge (dpc), except for 5- and 6-dpc. SAV3 RNA was detected in water samples by RT-qPCR between 7-and 16-dpc, with no detection after 16 dpc. For each individual fish, the viral load in the heart at 35 dpc is indicated by a corresponding Ct value; absence of a Ct value denotes non-detection. The y-axis represents calculated copy numbers, divided into three sections (0-500, 500-15,000, and 15,000-80,0000) to highlight differences in shedding magnitude. The x-axis shows time after challenge (2-19 dpc). Data for NV-C, Cl-C, and Mi-C are shown in black, blue, and red, respectively.

**Figure 7.**
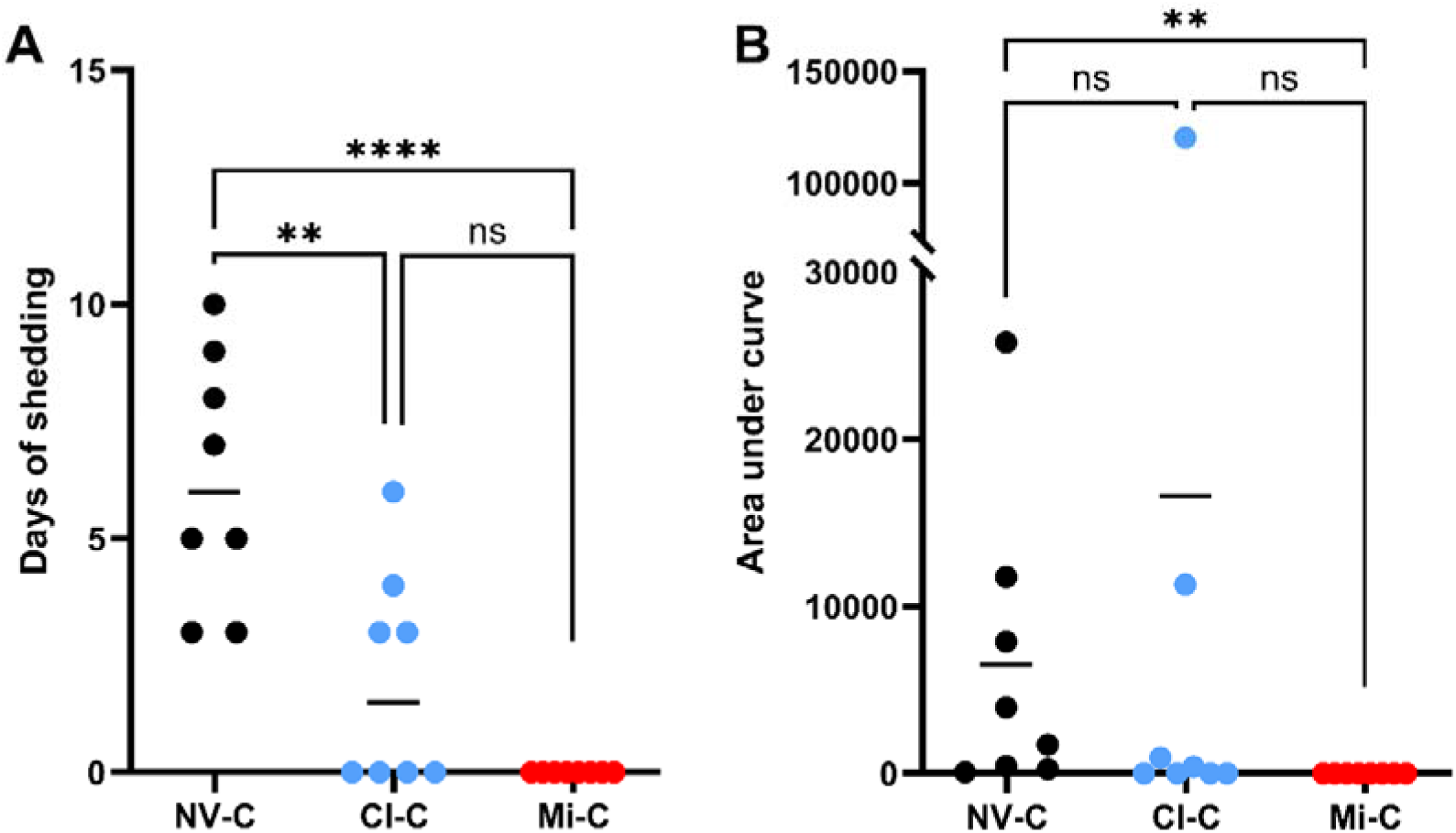
The duration of shedding and the cumulative viral shedding in SAV3 challenged individually housed fish. (A) The duration of shedding, calculated as the number of days shedding was detected, in Figure 6. Statistical significance was determined using Tukey’s multiple comparison testing and is indicated as follows: ns (not significant; ** (*p* < 0.01), **** (*p* < 0.0001). (B) The cumulative viral shedding over the experimental period, quantified as the area under the curve (AUC) for each fish (subfigure) in Figure 6. Statistical significance was determined using Kruskal-Wallis testing and is indicated as follows: ns (not significant), ** (*p* < 0.01). Black points represent non-vaccinated (NV-C), blue points represent Clynav (Cl-C) and red points represent AlphaJect Micro1-PD (Mi-C). Horizontal bars indicate group mean values.

Cumulative shedding over the experimental period was calculated as the area under the curve (AUC) for each individual tank (**Figure 6**). The cumulative shedding of Mi-C was significantly lower than in the NV-C (Kruskal-Wallis test, p=0.0013), (**Figure 7B)**. The Cl-C group included one high shedding individual, and the cumulative shedding in the Cl-C group did not differ significantly from the other groups. Across the 24 individual fish from the three vaccination groups, there was a very strong, positive correlation (Spearman r = 0.8289, p<0.0001) between cumulative shedding over the experimental period and viral load in the heart at 35 dpc (**Figure 8**). Given the log-log representation of both axes (Ct values) (**Figure 8**), the relationship between cumulative shedding (y) and viral load in the heart at 35 dpc, can be approximated by the power function y = x^0.9238^ + 2^0.8142^ = x^0.9238^ + 1.7626. This reflects a near-proportional scaling of cumulative shedding with the viral load in the heart, adjusted for the base-2 logarithmic nature of Ct values.

**Figure 8.**
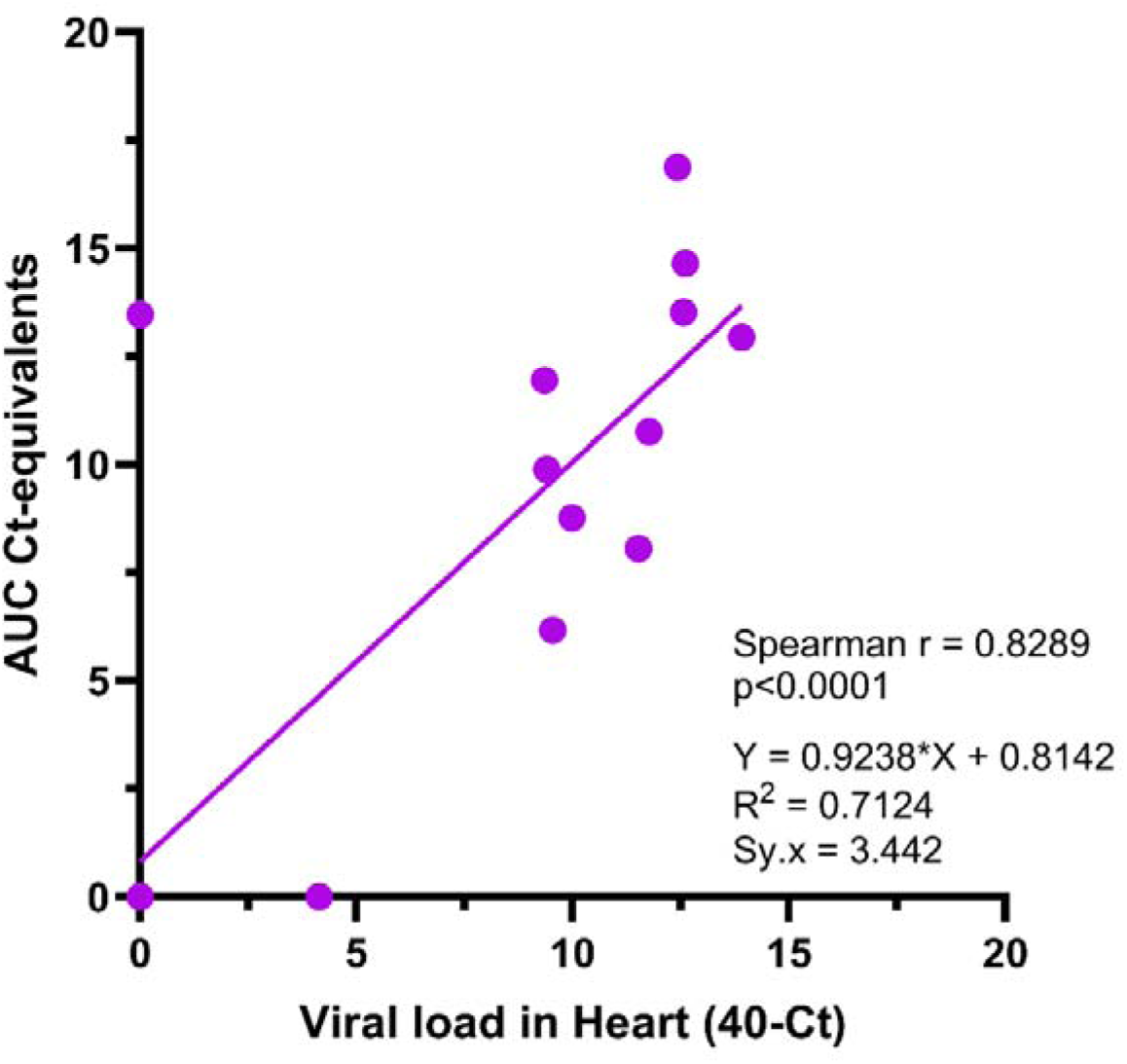
The relationship between cumulative viral shedding and viral load in the heart of SAV3-challenged, individually housed fish. A scatter plot illustrating the correlation between cumulative shedding (AUC, y-axis) and the corresponding viral load in heart at 35 dpc (x-axis) for the 24 individually housed fish (i.e. including non-vaccinated and Micro1-PD- and Clynav-vaccinated groups). Cumulative viral shedding over the experimental period was quantified as the area under the curve (AUC) for each fish (see subfigures in Figure 6) and log_2_-transformed to AUC ‘Ct-equivalents’. The plot includes results from a Spearman rank correlation test (Spearman r = 0.8289), and a linear regression (Y = 0.9238x + 0.8142). Note that the data point at (x,y) = (0,0) represents 13 double-negative individual fish.

In the cohort experiment, the kinetics of SAV3 shedding was monitored by analysis of water samples collected between 2- and 32 dpc. Virus shedding was detected in all cohort tanks with challenged fish (NV-C, Cl-C and Mi-C) (**Figure 9**, **Supplementary Figure 6**), but not in tanks with non-challenged fish (NV-NC, Cl-NC and Mi-NC, data not shown). Virus shedding was first detected at 4 dpc in NV-C and Cl-C tanks, and at 5 dpc in Mi-C tanks. In the NV-C and Cl-C tanks, shedding increased markedly after initial detection, peaking at 7-9 dpc, before declining and becoming sporadically detectable after ∼18 dpc (**Supplementary Figure 6**). In contrast, shedding in the Mi-C tanks increased after initial detection but did not exhibit a similarly clear early peak. Instead, viral RNA was consistently detected at relatively lower levels until the end of the experiment at 32 dpc (**Supplementary Figure 6**). The sporadic detection of SAV3 RNA in NV-C and Cl-C tanks after 18 dpc suggests that shedding levels dropped to near the limit of detection, resulting in intermittent positive samples. The duration of viral shedding, as measured by the number of days shedding was detected, was significantly higher in the Mi-C group than in both the NV-C (Tukey’s multiple comparison test, p=0.0012) and Cl-C (p<0.0001) groups (**Figure 10A**). The cumulative shedding over the experimental period was calculated as the area under the curve (AUC) for each challenged cohort tank in (**Figure 9**). The cumulative shedding of the Mi-C group was significantly lower than in the NV-C group (Kruskal-Wallis test, p=0.0181) **(Figure 10B**).

**Figure 9.**
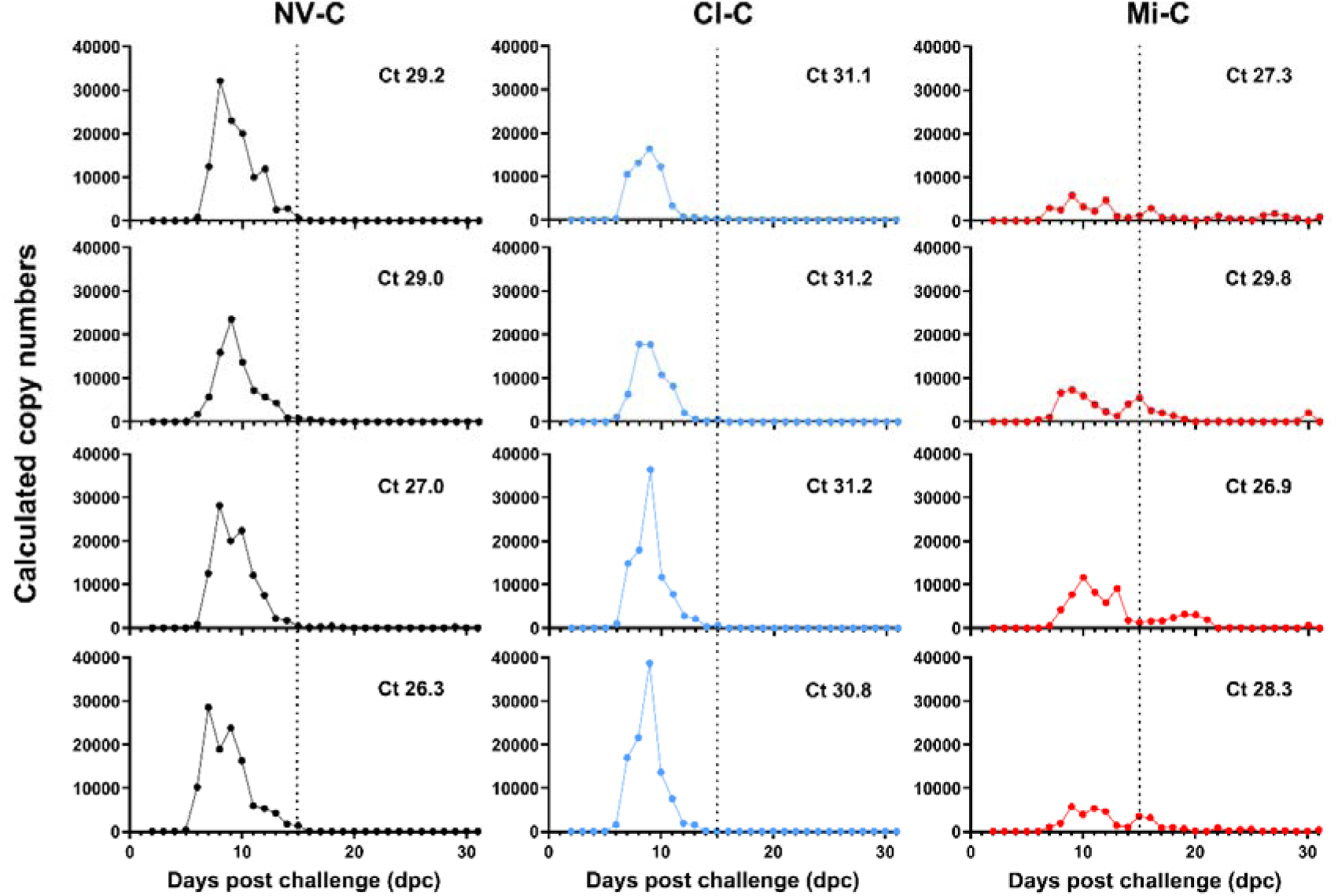
Temporal virus shedding profiles in fish cohorts after SAV3 challenge. Between 2 and 32 days post-challenge (dpc), water samples (100 mL) were collected daily from four replicate tanks per vaccination group (Non-vaccinated, NV-C; Clynav, Cl-C and AlphaJect Micro1-PD, Mi-C), each containing challenged fish. SAV3 RNA was detected in water samples by RT-qPCR from 4 dpc. For each tank, the mean viral load in heart tissue of 8 fish sampled at 32 dpc is indicated by a corresponding Ct value. The y-axis represents calculated copy numbers, and the x-axis shows days post challenge (dpc).

**Figure 10.**
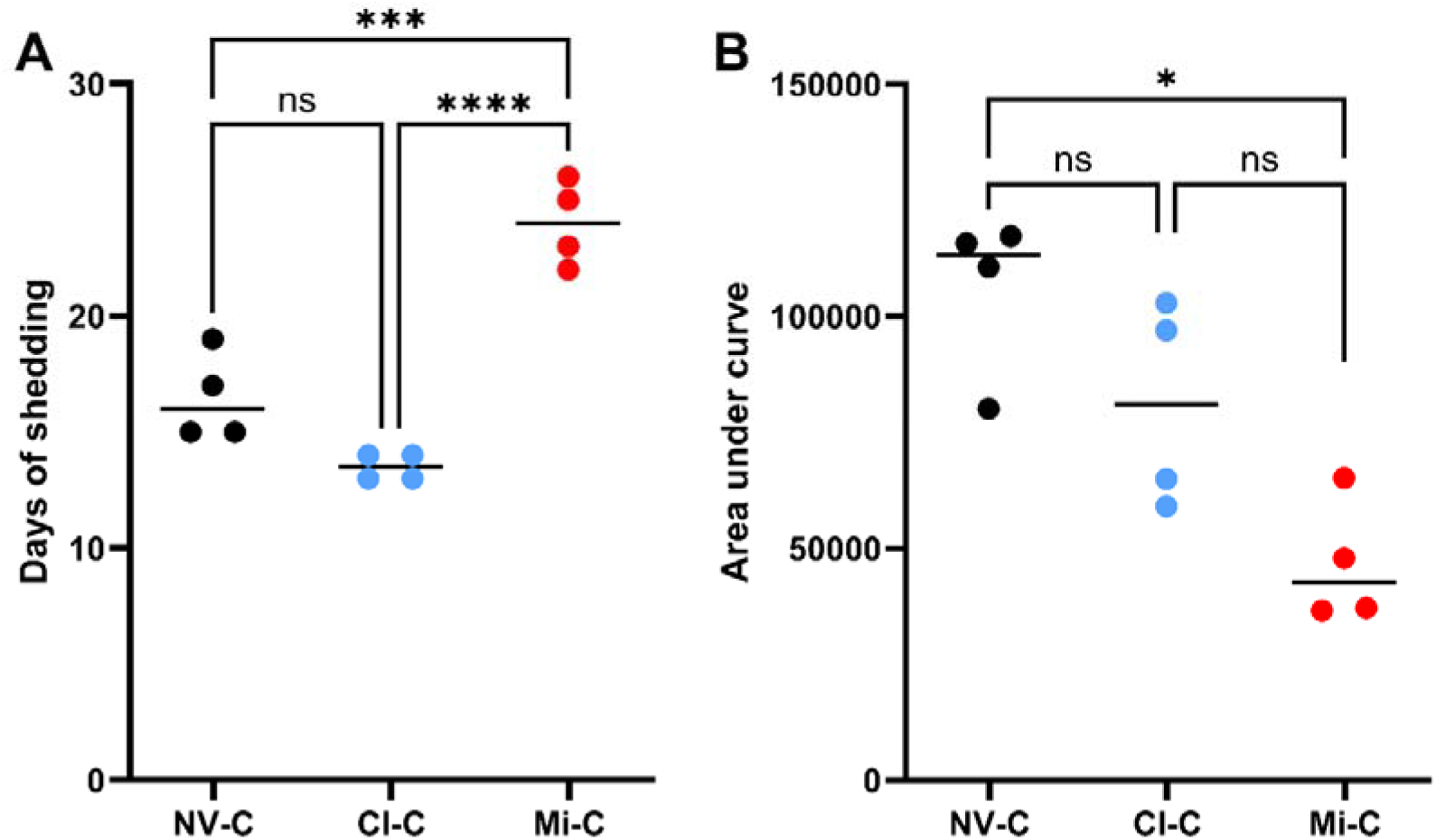
The duration of shedding and the cumulative viral shedding in SAV3 challenged fish cohorts. (A) The duration of shedding, calculated as the number of days shedding was detected, in Figure 9. Statistical significance was determined using Tukey’s multiple comparison testing and is indicated as follows: ns (not significant; *** (*p* < 0.001), **** (*p* < 0.0001). (B) The cumulative viral shedding over the experimental period was quantified as the area under the curve (AUC) for each tank (subfigure) in Figure 9. Statistical significance was determined using Kruskal-Wallis testing and is indicated as follows: ns (not significant), * (*p* < 0.05). Black points represent non-vaccinated (NV-C), blue points represent Clynav (Cl-C) and red points represent AlphaJect Micro1-PD (Mi-C). Horizontal bars indicate group mean values.

### 3.4 Viral load in tissues at experiment termination

The viral load of SAV3 was examined in individually housed, challenged fish at 35 dpc by analysing heart, pancreas and spleen tissue samples by RT-qPCR (**Figure 11**). In the heart samples, all the eight non-vaccinated (NV-C) fish were positive for SAV3 compared to three of eight Clynav-vaccinated (Cl-C) and one of eight Micro1-PD-vaccinated (Mi-C) fish. By Dunn’s multiple comparison test, the viral load in the NV-C group was significantly higher than in the Cl-C group (p=0.0098) and the MI-C group (p=0.0003). In pancreas, three NV-C fish and one Cl-C fish were SAV3 positive, while no Mi-C fish were positive. In spleen, seven NV-C fish were SAV3 positive, compared to four fish in both the Cl-C and Mi-C groups. No significant differences in viral load were observed in the pancreas or spleen by Dunn’s multiple comparison test.

**Figure 11.**
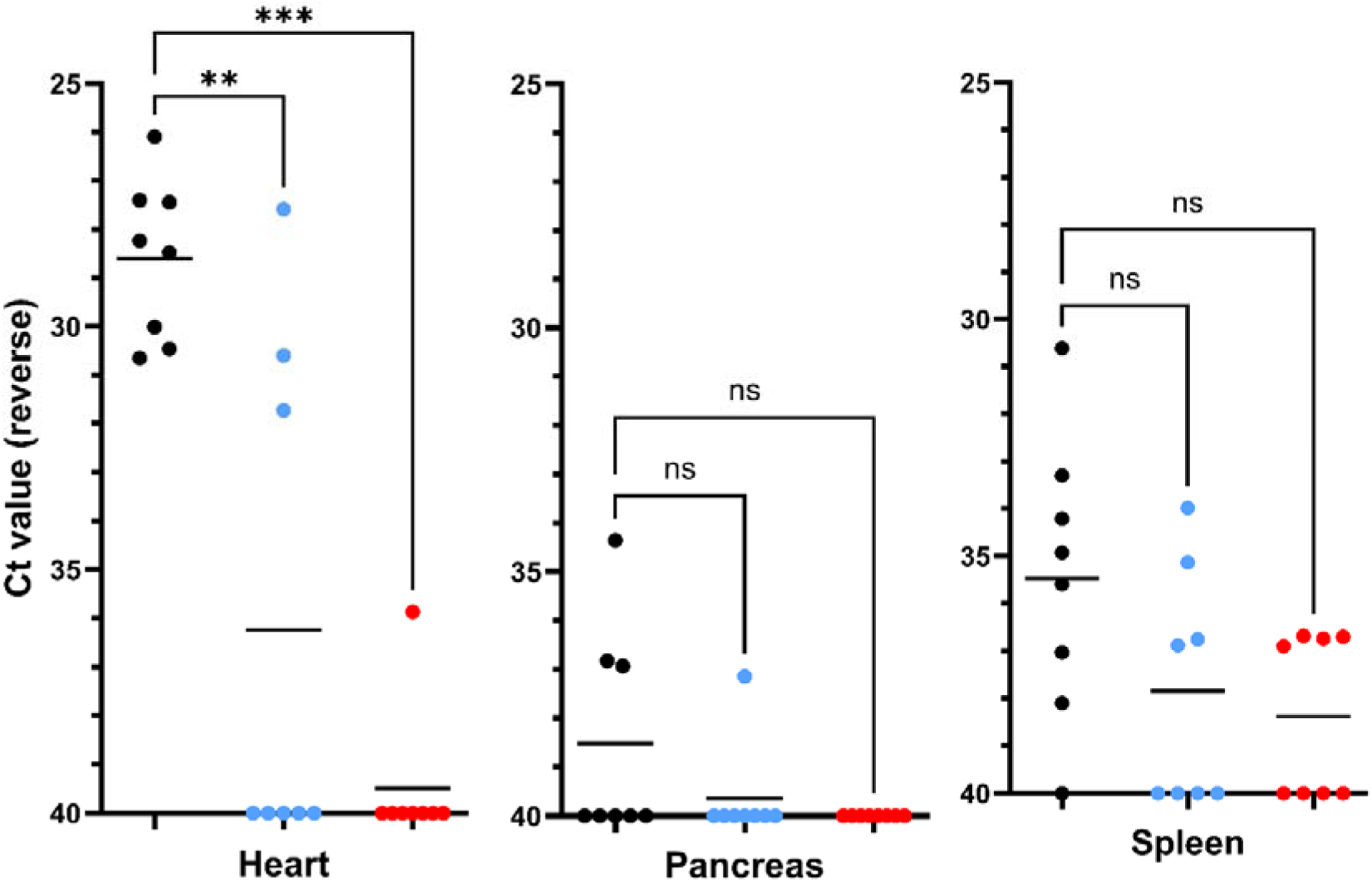
Viral load in heart, pancreas and spleen of challenged, individually housed fish at 35 dpc. Tissue samples were collected from eight individual fish per vaccination group (non-vaccinated (NV, black), Clynav-vaccinated (Cl, blue) and AlphaJect Micro1-PD-vaccinated (Mi, red)) at 35 days post-challenge (dpc). Viral RNA (SAV3) was quantified by RT-qPCR, with results presented as reverse Ct values (y-axis). Asterisks (* *p* < 0.05; ** *p* < 0.01; *** *p* < 0.001; ns = not significant) indicate statistical significance based on Dunn’s multiple comparison testing of vaccinated groups against the non-vaccinated group. Horizontal bars show group mean Ct values.

In the cohort experiment, the viral load of SAV3 was examined in the heart, pancreas and spleen samples collected from eight fish per cohort tank at 32 dpc. Viral RNA was not detected in any of the non-challenged groups, regardless of tissue (data not shown). In contrast, SAV3 was demonstrated in most challenged fish, and the data were analysed by two-way ANOVA to assess the effects of tank (replicate) and vaccination status (NV, Clynav, Micro1-PD) (**Figure 12**). In the heart, there was no significant tank effect (p=0.49), but a significant effect of vaccination (p=0.0034). By Tukey’s multiple comparison testing, the viral load in the Clynav group was significantly lower than in the NV group (p=0.0069) and the Micro1-PD group (p=0.0124). In the pancreas, there was a trend towards a tank effect (p=0.0817), but a significant effect of vaccination (p=0.0016). Tukey’s multiple comparison test revealed that the viral load in the Micro1-PD group was significantly higher than in the NV group (0.0064) and the Clynav group (p=0.0041). In the spleen, there was no significant tank effect (p=0.12), but a significant effect of vaccination (p=0.0201). Tukey’s multiple comparison test showed that the viral load in the Micro1-PD group was significantly higher than in the NV group (p=0.0462) and the Clynav group (p=0.0349).

**Figure 12.**
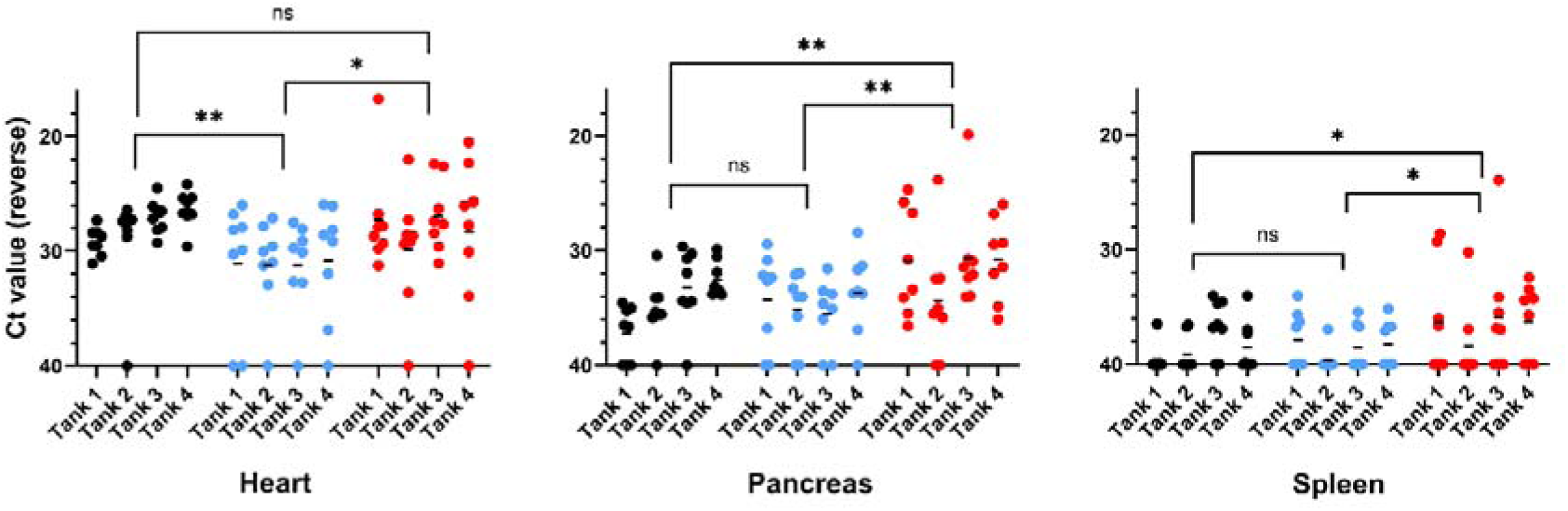
Viral load in heart, pancreas and spleen of SAV3 challenged cohort fish sampled at 32 dpc. For each vaccination group (non-vaccinated (black), Clynav-vaccinated (blue) and AlphaJect Micro1-PD-vaccinated (red)), eight fish were sampled from each of four replicate tanks (tank 1-4) at 32 days post-challenge (dpc). Viral RNA (SAV3) was quantified by RT-qPCR, with results presented as reverse Ct values (y-axis). Asterisks (ns = not significant; * *p* < 0.05; ** *p* < 0.01; *** *p* < 0.001) indicate statistically significant differences between vaccination groups based on a Tukey’s multiple comparison test. Horizontal bars represent the mean Ct values per tank.

### 3.5 SAV3 related histopathology in tissues at experiment termination

Histopathological lesions associated with SAV3 infection, were examined and scored in the heart, pancreas, and skeletal muscle of individually housed, challenged fish sampled at 35 dpc (**Figure 13**). In the heart, lesion scores for epicarditis, myocarditis and degeneration were generally low and showed no significant differences between vaccination groups (Dunn’s multiple comparison test). In the pancreas, all eight non-vaccinated fish exhibited lesions, compared to two of eight Clynav-vaccinated, and five of eight Micro1-PD-vaccinated fish. Lesion scores in non-vaccinated fish were significantly higher than in Clynav-vaccinated fish (p=0.0015) but not significantly different from those in Micro1-PD-vaccinated fish (p=0.0745). In red muscle, all eight non-vaccinated fish showed lesions, compared to four of eight Clynav-vaccinated, and one of eight Micro1-PD-vaccinated fish. Lesion scores in non-vaccinated fish were significantly higher than in both Clynav-vaccinated fish (p=0.0386) and Micro1-PD-vaccinated fish (p=0.0006). In white muscle, all eight non-vaccinated fish showed lesions, compared to three of eight Clynav-vaccinated, and two of eight Micro1-PD-vaccinated fish. Lesion scores in non-vaccinated fish were significantly higher than in both Clynav-vaccinated fish (p=0.0231) and Micro1-PD-vaccinated fish (p=0.0019).

**Figure 13.**
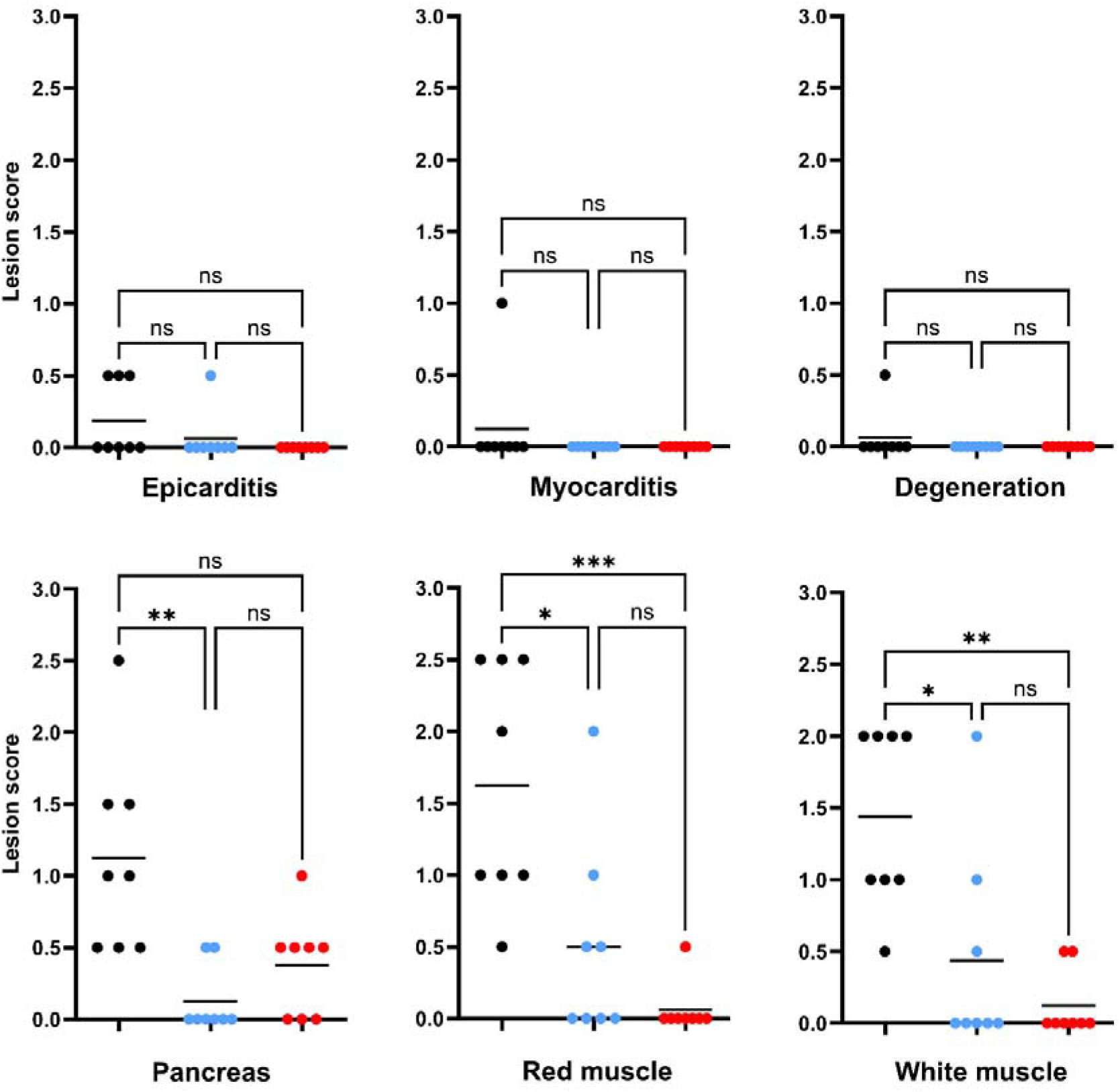
Histopathological lesion scores in heart, pancreas and skeletal muscle of challenged, individually housed fish at 35 dpc. Tissue samples were collected from the eight individual fish in each vaccination group (non-vaccinated (black), Clynav-vaccinated (blue) and AlphaJect Micro1-PD-vaccinated (red)) at 35 days post-challenge (dpc). Histopathological lesion types in the heart (epicarditis, myocarditis, degeneration) and lesions in pancreas, red muscle, and white muscle were scored from 0 (no lesions) to 3 (severe lesions). Statistical significance was determined using Dunn’s multiple comparison test among the challenged groups. Asterisks indicate levels of significance (* *p* < 0.05; ** *p* < 0.01; *** *p* < 0.001; ns = not significant). Horizontal bars indicate group means for lesion score.

In the cohort experiment, histopathological lesions associated with SAV3 infection and/or vaccination were examined and scored in the heart, pancreas, and skeletal muscle of fish sampled at 32 dpc (**Figure 14**). Among the non-challenged vaccination groups (NV-NC, Cl-NC, Mi-NC), myocarditis and degeneration lesions in the heart were absent, as were lesions in red and white muscle. Mild epicarditis was observed in two of eight Cl-NC fish, compared to one of eight Mi-NC fish and none of the NV-NC fish. Mild lesions in the pancreas, compatible with vaccination-induced lesions, were observed in three of eight Mi-NC fish. No pancreas lesions were observed in NV-NC or Cl-NC fish. Dunn’s multiple comparison test revealed no significant differences in lesion scores among the non-challenged groups, irrespective of tissue and lesion type. In the challenged vaccination groups (NV-C, Cl-C, Mi-C), few specimens exhibited lesions in the heart, although four of eight NV-C fish had predominantly mild myocarditis scores. In the pancreas, seven of eight NV-C fish exhibited lesions (mean lesion score = 1.63), compared to five of eight Cl-C fish (mean lesion score = 1.00) and seven of eight Mi-C fish (mean lesion score = 1.38). For red muscle, lesions were observed in seven of eight NV-C fish (mean lesion score = 1.50), compared to six of eight Cl-C fish (mean lesion score = 1.00) and four of seven Mi-C fish (mean lesion score = 0.71). In white muscle, six of eight NV-C fish had lesions (mean lesion score = 0.81), compared to six of eight Cl-C fish (mean lesion score = 0.75) and one of seven Mi-C fish (mean lesion score = 0.14). As with the non-challenged groups, Dunn’s multiple comparison test revealed no significant differences in lesion scores among the challenged cohort groups, irrespective of tissue and lesion type.

**Figure 14.**
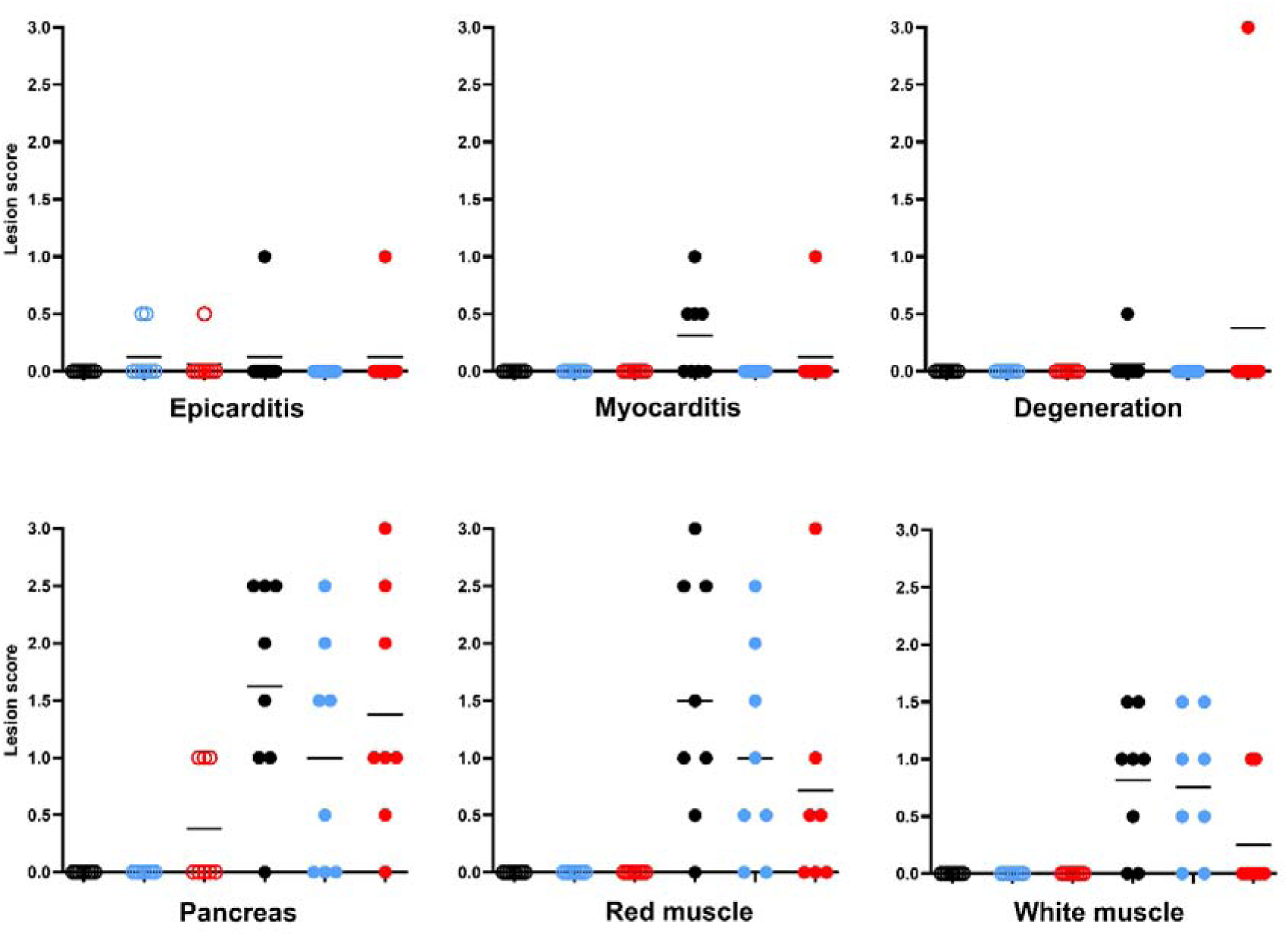
Histopathological lesion scores in heart, pancreas and skeletal muscle of cohort fish sampled at 32 dpc. Tissue samples were collected from eight individual fish in each vaccination group (non-vaccinated (black), Clynav-vaccinated (blue) and AlphaJect Micro1-PD-vaccinated (red)) at 32 dpc. Histopathological lesion types in the heart (epicarditis, myocarditis, degeneration) and lesions in pancreas, red muscle, and white muscle were scored from 0 (no lesions) to 3 (severe lesions). Open circles represent non-challenged fish, and filled circles represent challenged fish. Horizontal bars indicate group means for lesion score. Statistical significance was determined using Dunn’s multiple comparison test, comparing lesion scores among the three non-challenged groups and among the three challenged groups, respectively. No significant differences were found.

## 4. Discussion

In this study, we assessed the efficacy of two commercial SAV3 vaccines in reducing viral shedding in Atlantic salmon post-smolts following an experimental bath challenge. Fish from each treatment group were challenged separately, and then distributed between cohort and individual tanks. To monitor SAV3 shedding, water samples were collected daily and analysed for viral RNA using RT-qPCR. During the experiment, all fish in the cohort tanks developed clinicals signs (e.g. fin rot) subsequently attributed to *Tenacibaculum dicentrarchi* infection. By contrast, no fish in the individual tanks exhibited such signs. Between the individually housed fish, both the Micro 1-PD- and Clynav-vaccinated groups demonstrated significant reductions in shedding as compared to non-vaccinated controls, with the Micro 1-PD group showing the most pronounced effect. However, the difference in shedding between the two vaccinated groups was not statistically significant. These trends were mirrored in weight data, viral load in the heart at 35 dpc, and, to a lesser extent, histopathology scores in the pancreas and muscle at 35 dpc. In cohort tanks, shedding kinetics differed from those observed in individual tanks. Micro 1-PD-vaccinated cohorts exhibited significantly lower cumulative shedding than non-vaccinated controls but demonstrated a significantly prolonged shedding duration. Clynav-vaccinated cohorts, meanwhile, showed a shorter shedding duration and reduced cumulative shedding compared to NV-C cohorts, though these differences did not reach statistical significance. These patterns aligned with viral loads in heart, pancreas and spleen at 32 dpc, but were not reflected in weight data or histopathology scores.

The shedding patterns observed in challenged non-vaccinated fish, in both cohort and individual tanks, align with the findings by Andersen and coworkers (Andersen, Hodneland, and Nylund 2010), who detected SAV3 RNA shedding between 4 and 13 days post infection (dpi) following an intraperitoneal challenge with SAV3, with a peak at 6 dpi. In the present study, shedding was assessed by quantifying SAV3 RNA as a proxy for infectious virions.

However, fish develop neutralizing antibodies (Abs) within approximately two weeks following SAV3 infection (McLoughlin et al. 1996), as well as after vaccination with inactivated whole virus (Xu, Mutoloki, and Evensen 2012) or DNA (Thorarinsson et al. 2021). Consequently, the gradual development of neutralizing Abs in challenged non-vaccinated fish, and remaining pre-existing vaccine-induced neutralizing Abs in vaccinated groups, likely influenced the shedding of infectious SAV3 virions in ways not fully captured by RT-qPCR-based analysis. While both vaccines reduced SAV3 RNA shedding during the first 15-18 dpc, they may have conferred an even greater reduction in the shedding of infectious virions. To address this uncertainty, future studies of vaccine mediated shedding-reduction should incorporate functional infectivity assays of shed virus. Thorarinsson and coworkers, using a cohabitation challenge approach, demonstrated that the ‘infection pressure’ from vaccinated fish was higher in Micro 1-PD-vaccinated groups than in Clynav-vaccinated groups (Thorarinsson et al. 2021). However, in their study, cohabitation was initiated 21 days after the vaccinated fish had been challenged - a timeframe that, based on the data presented here, suggests that the bulk of viral shedding had likely already occurred.

Except for the Micro 1-PD-vaccinated cohorts, SAV3 RNA shedding declined to non-detectable or sporadic levels from approximately 15 – 18 dpc in all cohorts and individual tanks. This timing coincides with the expected emergence of both neutralizing and non-neutralizing Abs in infected fish (McLoughlin et al. 1996). In individual tanks, fewer vaccinated fish shed detectable levels of SAV3 RNA during the first 15–18 dpc, and in the cohorts, the area under the curve (AUC) for cumulative shedding was lower for vaccinated tanks. These findings may suggest that pre-existing Abs in vaccinated fish were able to reduce SAV3 RNA shedding while a secondary memory response was being mounted in response to the infection.

The observed difference in SAV3 RNA shedding between cohort and individual tanks for the Micro 1-PD vaccine is interesting. Data from the individual tanks indicate that Micro 1-PD was superior to Clynav in both reducing shedding and viral load in the heart. In contrast, under the influence of bacterial co-infection and/or the relative more crowded environment, Micro 1-PD cohort tanks exhibited significantly longer SAV3 shedding than even the non-vaccinated tanks. Based on the above reasoning regarding the role of antibodies, this breakdown in shedding-reducing capacity may be attributed to a hypothetically impaired ability to mount or maintain sufficient Ab levels. The significantly higher viral load in heart, pancreas and spleen at 32 dpc, could be the effect of potentially lower neutralizing Ab levels, or alternatively, may be the source of virus for the shedding levels observed in these cohorts.

When evaluating fish vaccines, efficacy is typically assessed using parameters such as morbidity, pathogen load, time to illness or death, histopathological damage, mortality, and transmission to naïve hosts (Midtlyng 2016). The use of high-throughput water sampling and RT-qPCR, as demonstrated in this study, may facilitate the inclusion of viral shedding as a standard efficacy parameter in future vaccine evaluations. A vaccine’s ability to reduce shedding, and thus lower the environmental pathogen load, is of fundamental importance for achieving herd immunity or resilience (Doeschl-Wilson et al. 2021). So-called ‘leaky’ vaccines protect individual fish against severe disease without adequately reducing shedding of the pathogen. By promoting healthy shedders and potentially driving evolution of pathogen virulence, leaky vaccines may compromise protection despite being efficacious in reducing disease severity in individuals (Read et al. 2015). Given that vaccination of farmed fish often results in highly variable individual vaccine responses (own observations), the presence of healthy super-shedders can contribute to undermining herd immunity and negatively impact overall disease resilience.

Using a high-throughput high-sensitivity method, developed for water sampling and RT-qPCR analysis, we detected SAV3 shedding from individually housed fish. At the individual level, we demonstrated a strong association between SAV3 shedding and RT-qPCR positivity in heart tissue at 35 dpc. Quantitatively, this relationship is characterized by a near-linear correlation on a log-log scale between the cumulative viral shedding (expressed as virus copies) and the load of virus copies in heart tissue at 35 dpc, approximated by the power function y = x^0.9238^ + 1.7626. Though more data is needed, the latter cautiously suggests a near 1:1 scaling between viral load in the heart and cumulative shedding. The ability to non-lethally monitor individual fish over time provides a powerful tool for investigating inter-individual variability in response to treatments such as vaccination, and for elucidating the genetic factors that influence vaccine efficacy and disease resistance.

In conclusion, the present study demonstrates that both Micro1-PD and Clynav vaccines significantly reduce SAV3 shedding in individually housed Atlantic salmon, with Micro1-PD exhibiting a more pronounced effect. However, under cohort conditions complicated by bacterial co-infection, the efficacy of Micro1-PD in reducing shedding duration was compromised, suggesting that vaccine efficacy can be context-dependent. Our findings underscore the utility of high-throughput water sampling and RT-qPCR as a non-lethal approach for monitoring presence and shedding of viral pathogens. Future studies should incorporate functional infectivity assays to distinguish between viral RNA and infectious virions, and further explore the immunological mechanisms underlying the observed differences in shedding dynamics, particularly the role of neutralizing antibodies. Ultimately, reducing viral shedding is critical for achieving herd immunity and limiting disease transmission. Our results provide a foundation for integrating shedding data as a standard parameter in the evaluation of fish vaccines.

## Acknowledgements

The authors acknowledge Ivar-Helge Matre, Tone Knappskog and Lise Dyrhovden for their excellent work with rearing and vaccinating fish, as well as the essential involvement of Tuva Sagstad, Thomas Røttingen and Joachim Nordbø in the challenge experiments. The comprehensive water sampling was only possible due to the excellent help from: Stig Mæhle, Roger Lille-Langøy, Laila Unneland, Sofie Knutar, Maren Hoff Austgulen, Elisabeth Stöger, Håkon Berg-Rolness, and Johanne Øyro. The work was funded by the FHF – Norwegian Seafood Research Fund (grant 901796).

## Data Availability Statement

Data will be made available on request.

## Supplementary Figures

**Supplementary Figure 1.**
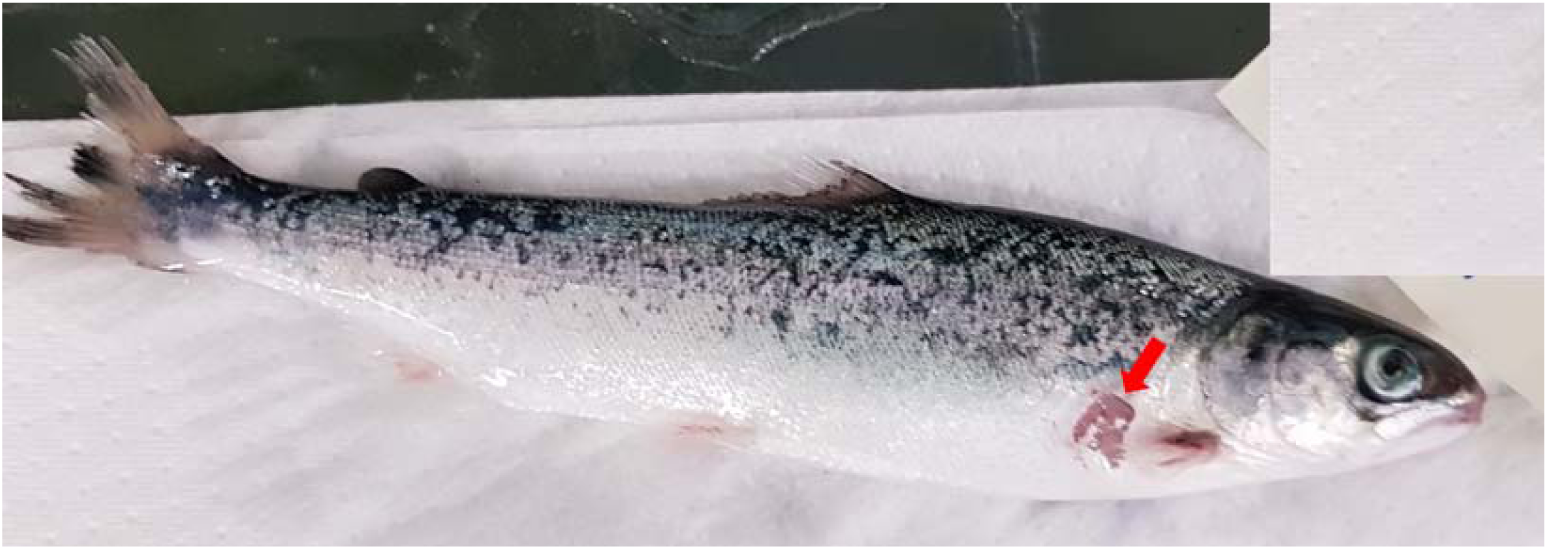
Euthanized fish with macro-clinical symptoms typical of the moribund fish in the cohort experiment. Euthanized moribund fish from a non-challenged control tank. Pectoral, dorsal and caudal fins exhibit marked fin erosion, Hemorrhages are seen at the base of anal and pelvic fins, and in the mouth region. Red arrow shows open ulcer.

**Supplementary Figure 2.**
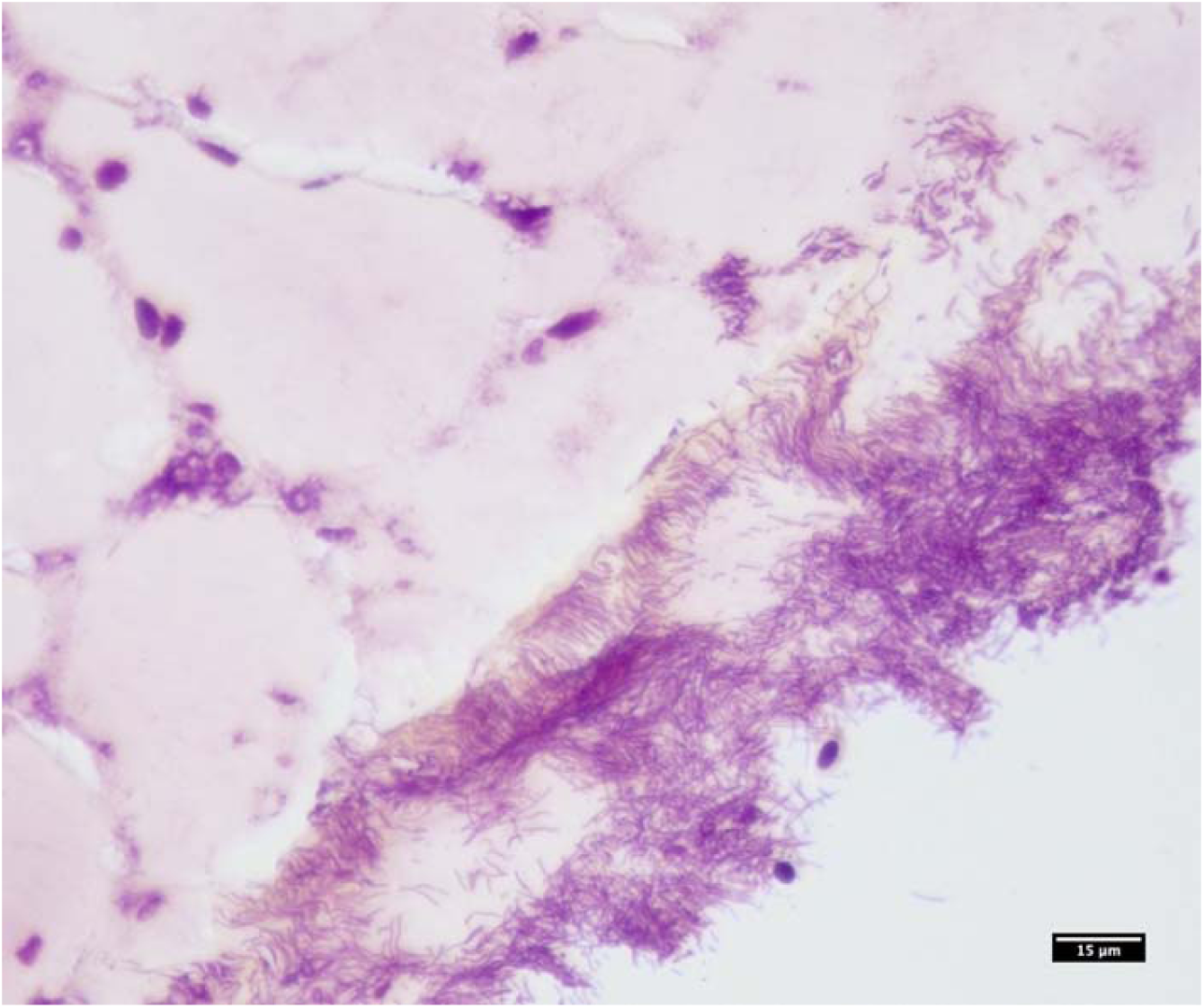
Micrograph of skin ulcer stained with modified Gram staining. Skin section from moribund fish stained with modified Gram staining shows abundant gram-negative rod-shaped, filamentous bacteria consistent with *Tenacibaculum* spp. Scale bar: 15 μm.

**Supplementary Figure 3.**
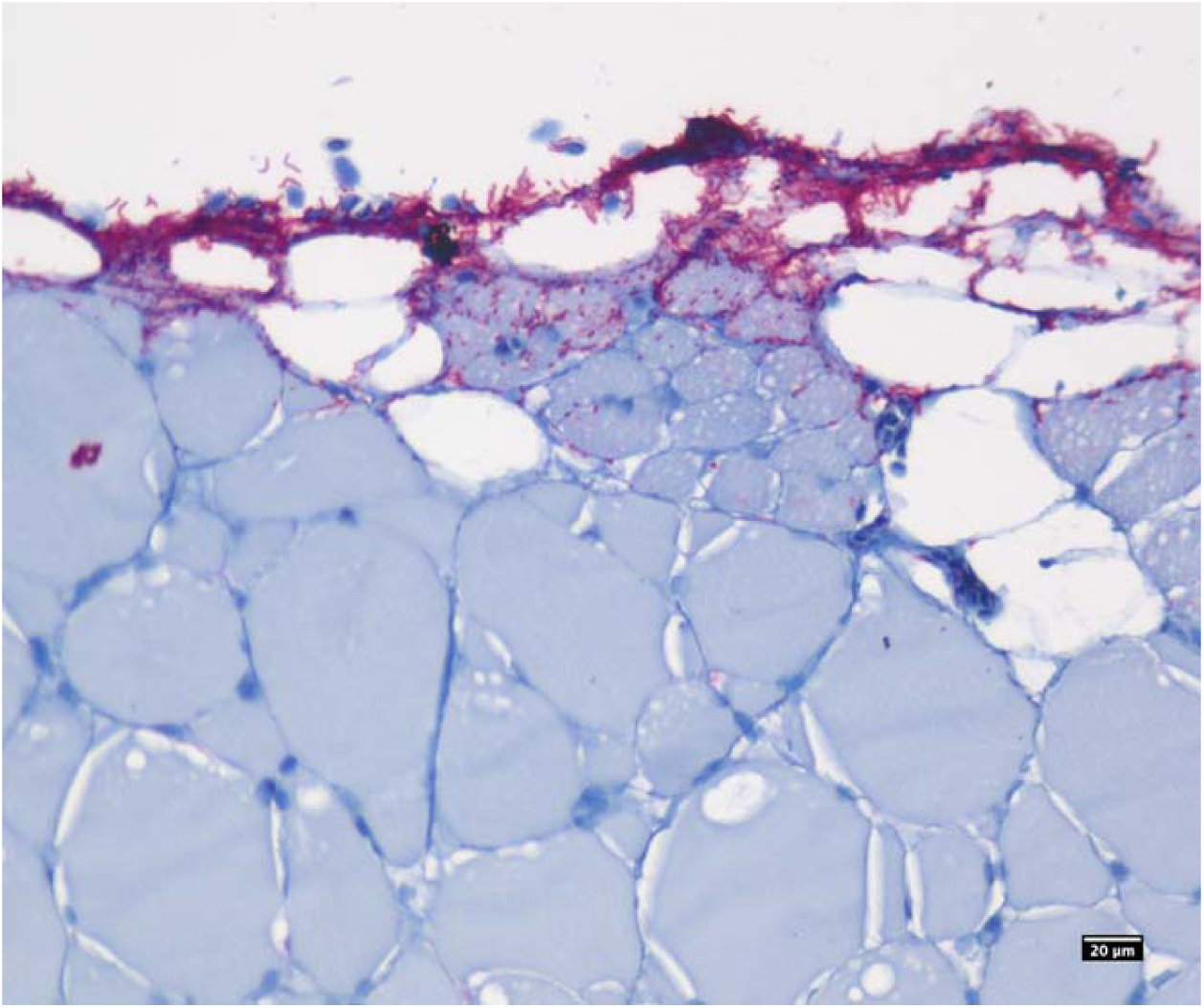
Micrograph of skin ulcer immunohistochemically stained with a polyclonal antiserum against *Tenacibaculum* spp. Indicated by red color, the staining demonstrates the abundant presence of *Tenacibaculum* spp. localized in the ulcerated upper layer of the skin, with some infiltration into the underlying muscle. Scale bar: 20 μm.

**Supplementary Figure 4.**
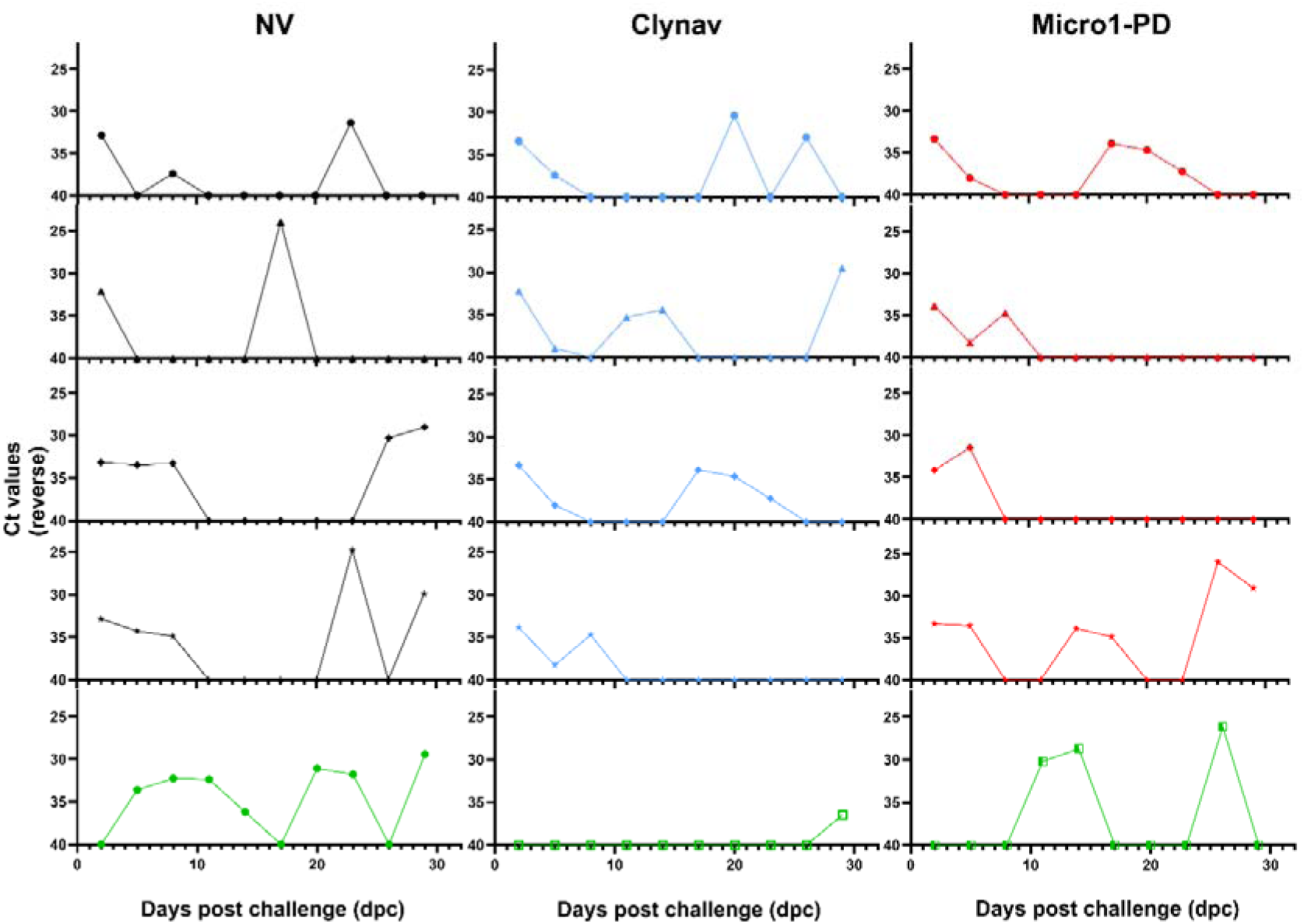
Profiles of temporal shedding of *Tenacibaculum dicentrarchi* in fish cohort groups. Water samples (100 mL) were collected daily from SAV3 challenged (four tanks per vaccination group) and non-challenged (one tank per vaccination group) groups between 2- and 32-days post-challenge (dpc). Using qPCR, presence of *Tenacibaculum dicentrarchi* was analyzed in water samples taken every third day from 2 dpc. For each vaccination group (NV in black, Clynav in blue and AlphaJect Micro1-PD in red), the four upper panels show the results from four challenged replicate tanks and the lower panel shows results from a non-challenged tank (green). The y-axis represents reverse Ct values, indicating the presence of bacterial DNA, and the x-axis represents days post challenge (dpc).

**Supplementary Figure 5.**
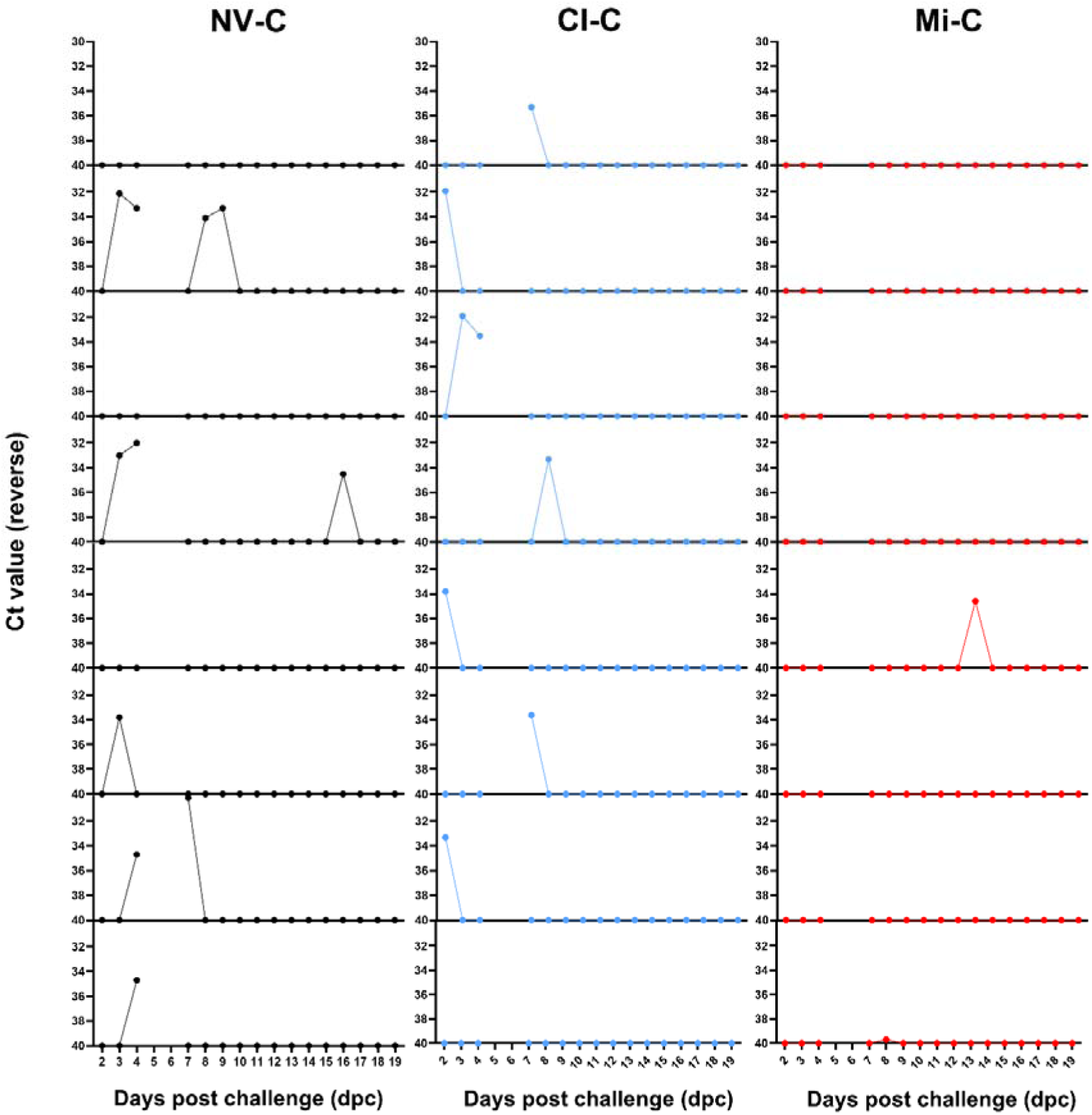
Profiles of temporal shedding of *Tenacibaculum dicentrarchi* in individual fish after SAV3 bath challenge. Water samples (100 mL) were collected daily from eight individual fish per challenged vaccination group (Non-vaccinated, NV-C; Clynav, Cl-C and AlphaJect Micro1-PD, Mi-C) between 2-and 35-days post-challenge (dpc), except for 5- and 6-dpc. Using qPCR, the presence of *Tenacibaculum dicentrarchi* was analyzed in water samples taken from 2 dpc. The y-axis represents reverse Ct values, indicating presence of bacterial DNA, and the x-axis represents days post challenge (dpc).

**Supplementary Figure 6.**
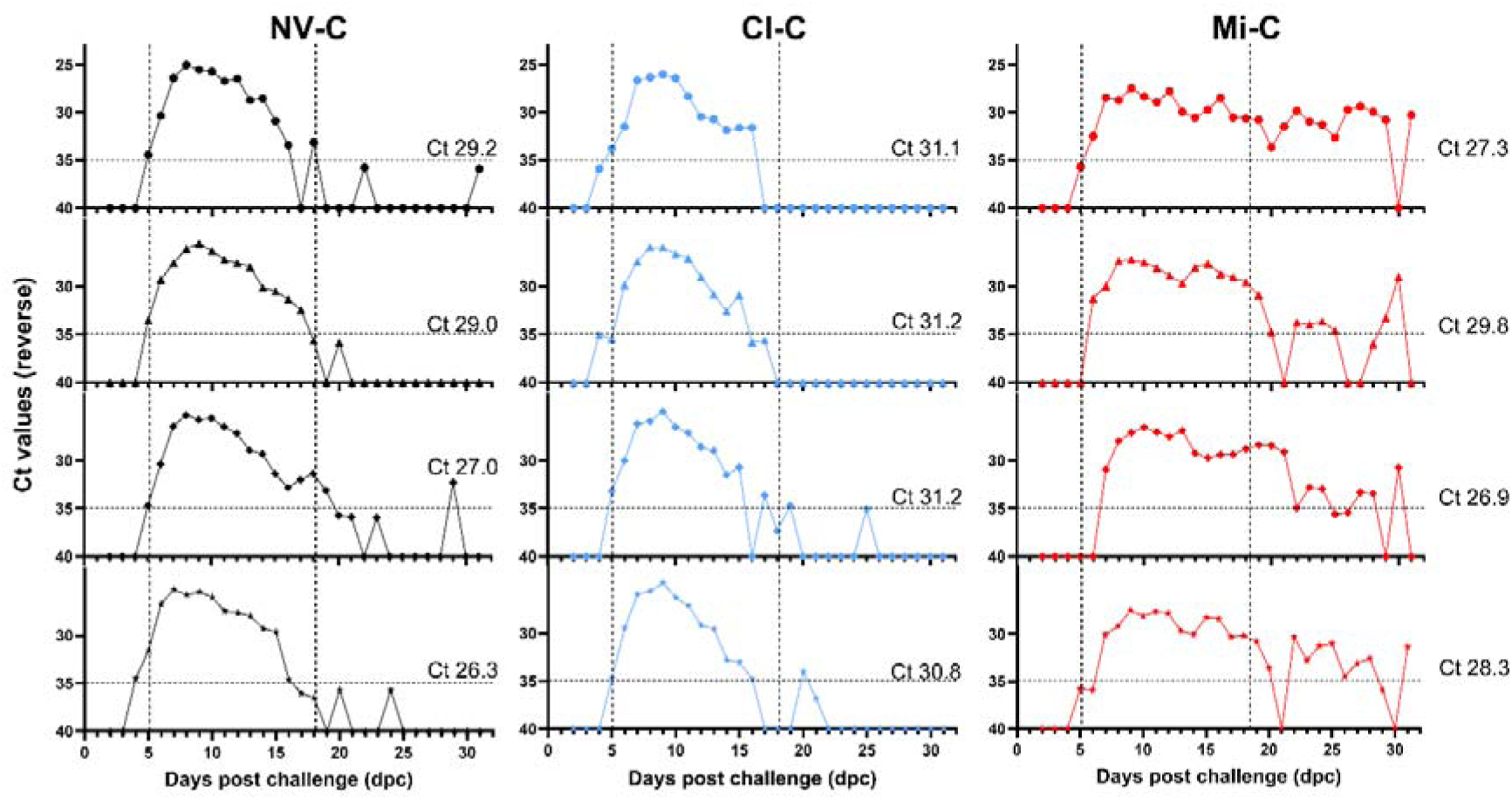
Temporal virus shedding profiles in fish cohorts after SAV3 challenge. Between 2 and 32 days post-challenge (dpc), water samples (100 mL) were collected daily from four replicate tanks per vaccination group (Non-vaccinated, NV-C; Clynav, Cl-C and AlphaJect Micro1-PD, Mi-C), each containing challenged fish. SAV3 RNA was detected in water samples by RT-qPCR from 4 dpc. For each tank, the mean viral load in heart tissue of 8 fish sampled at 32 dpc is indicated by a corresponding Ct value. The y-axis represents reverse Ct values, and the x-axis shows days post challenge (dpc).

---

**Supplementary Table 1.**
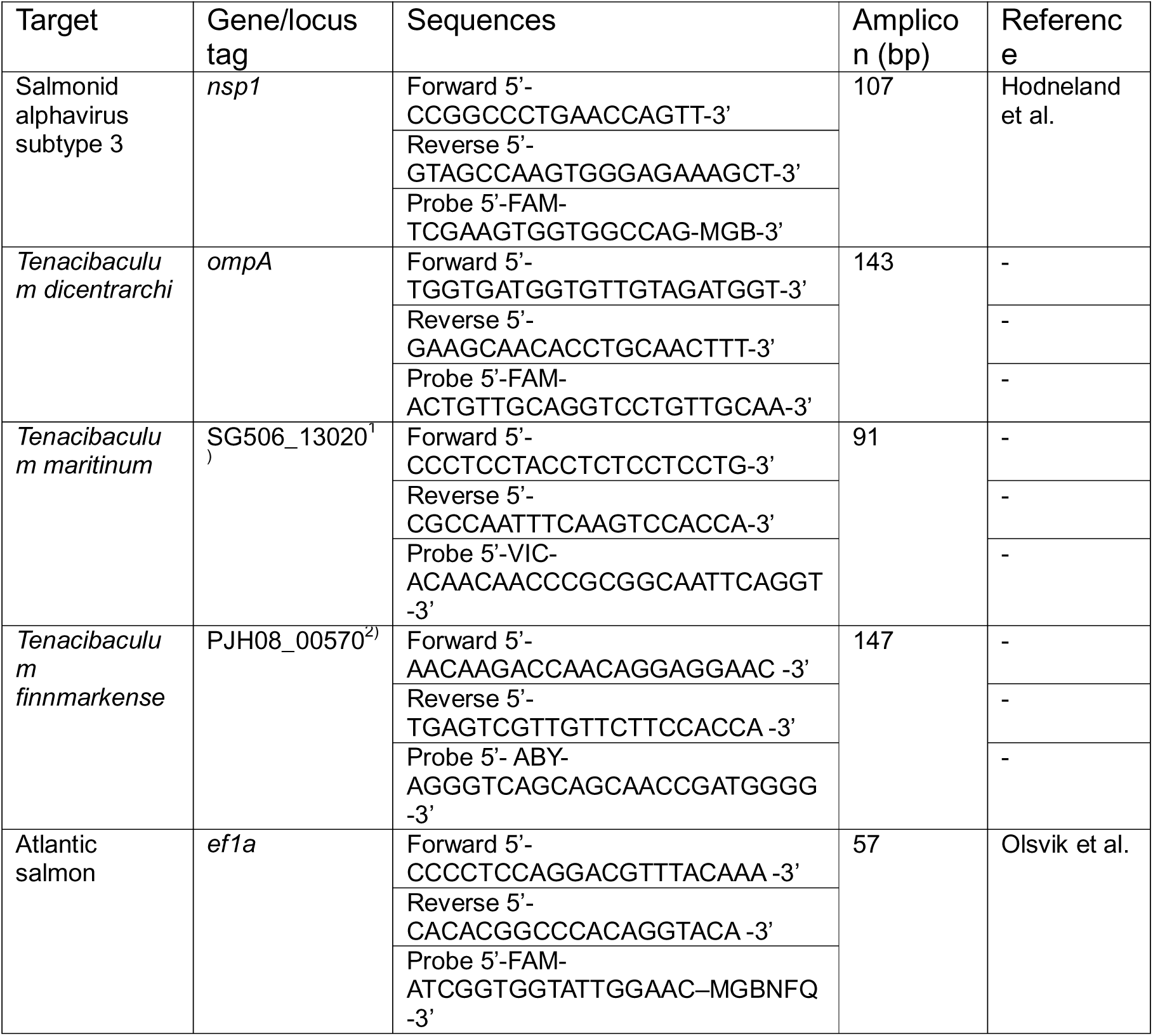
Primers and probes used in RT-qPCR and qPCR analyses. 1) DUF4114 domain-containing protein, GenBank: XKW65476.1 2) GEVED domain-containing protein, GenBank: WCC44957.1 Hodneland K, Endresen C. Sensitive and specific detection of Salmonid alphavirus using real-time PCR (TaqMan (R)). Journal of Virological Methods. 2006;131(2):184-92. Olsvik PA, Lie KK, Jordal A-EO, Nilsen TO, Hordvik I. Evaluation of potential reference genes in real-time RT-PCR studies of Atlantic salmon. BMC Molecular Biology. 2005;6(1):21.

**Supplementary Table 2.** Semi-quantitative system for scoring PD-related tissue lesions. Column 1 lists semi-quantitative scores ranging from 0 to 3, representing increasing severity. Column 2 is subdivided into three categories, epicarditis, myocarditis and degeneration, each detailing the histopathological changes in heart associated with the scores. Columns 3 and 4 describe the corresponding histopathological findings in the pancreas and skeletal muscle, respectively. The table outlines progressive pathological changes from normal tissue at S=0 to severe lesions at S=3, providing a standardized framework for assessing tissue damage.

| Score<br>(0-3) | Heart |  |  | Pancreas | Skeletal<br>muscle |
| --- | --- | --- | --- | --- | --- |
|  | Epicarditis | Myocarditis | Degeneration |  |  |
| S = 0 | Normal | Normal | Normal | Normal | Normal |
| 0 < S < 1 | Focal changes with limited number of mononuclear inflammatory cells (2-4 foci) infiltrating the epicardium | Minor inflammatory cell-infiltration of compactum | Focal myocardial degeneration (<50 fibres affected) | Focal necrosis of exocrine pancreatic cells (acinar cells); significant intact acinar cells | Focal necrosis of fibres ± inflammation |
| 1 ≤ S < 2 | Multifocal infiltration of inflammatory cells (>5 cell layers) infiltration the epicardium | Focal to multifocal inflammatory cell-infiltration (2-5 foci) of compactum | Multifocal myocardial degeneration (50-75 fibres affected) | Mild to moderate necrosis of exocrine pancreatic cells; some intact acinar cells | Mild to moderate necrosis of fibres ± inflammation |
| 2 ≤ S < 3 | High number of inflammatory cells (>10 cell layers) infiltration most of the epicardium | Multifocal to diffuse inflammatory cell-infiltration in compact or spongy layers | Multifocal to severe myocardial degeneration (75-100 fibres affected) | Moderate to severe necrosis of exocrine pancreatic cells; a few intact acinar cells | Moderate to severe necrosis of fibres ± inflammation |
| S = 3 | Severe inflammation (>15 cell layers) involving > 75% of the epicardium | Severe or widespread inflammatory cell-infiltration in compact or spongy layers in a diffuse distribution pattern | Severe to diffuse myocardial degeneration (>100 fibres affected) | Total absence of the exocrine pancreas | Diffuse or widespread necrosis of fibres ± inflammation |

